# Genetic paths to evolutionary rescue and the distribution of fitness effects along them

**DOI:** 10.1101/696260

**Authors:** Matthew M Osmond, Sarah P Otto, Guillaume Martin

**Author notes:** Center for Population Biology, University of California, Davis (Matthew M Osmond).

## Abstract

The past century has seen substantial theoretical and empirical progress on the genetic basis of adaptation. Over this same period a pressing need to prevent the evolution of drug resistance has uncovered much about the potential genetic basis of persistence in declining populations. However, we have little theory to predict and generalize how persistence – by sufficiently rapid adaptation – might be realized in this explicitly demographic scenario. Here we use Fisher’s geometric model with absolute fitness to begin a line of theoretical inquiry into the genetic basis of evolutionary rescue, focusing here on asexual populations that adapt through *de novo* mutations. We show how the dominant genetic path to rescue switches from a single mutation to multiple as mutation rates and the severity of the environmental change increase. In multi-step rescue, intermediate genotypes that themselves go extinct provide a ‘springboard’ to rescue genotypes. Comparing to a scenario where persistence is assured, our approach allows us to quantify how a race between evolution and extinction leads to a genetic basis of adaptation that is composed of fewer loci of larger effect. We hope this work brings awareness to the impact of demography on the genetic basis of adaptation.

---

Our understanding of the genetic basis of adaptation is rapidly improving due to the now widespread use of genomic sequencing (see examples in Bell 2009; Stapley *et al.* 2010; Dettman *et al.* 2012; Schlötterer *et al.* 2015). A recurrent observation, especially in experimental evolution with asexual microbes, is that the more novel the environment and the stronger the selection pressure, the more likely it is that adaptation primarily proceeds by fewer mutations of larger effect (i.e., that adaptation is oligogenic *sensu* Bell 2009). An extreme case is the evolution of drug resistance, which is often achieved by just one or two mutations (e.g., Bataillon *et al.* 2011; Pennings *et al.* 2014).

However, drugs, and other sufficiently novel environments, will often induce not only strong selection but also population decline. Such declines hinder both the production and maintenance of adaptive genetic variation (Otto and Whitlock 1997), thus impeding evolution and threatening extinction. Drug resistance evolution is a particular instance of the more general phenomenon of evolutionary rescue (Gomulkiewicz and Holt 1995; Bell 2017), where persistence requires sufficiently fast adaptive evolution.

Most theory on the genetics of adaptation (reviewed in Orr 2005) assumes constant population size and therefore does not capture the characteristic ‘race’ between adaptation and extinction that occurs during evolutionary rescue. Many models have been created to describe this race (reviewed in *Alexander et al.* 2014) but so far largely focus on two extreme genetic bases, both already introduced in Gomulkiewicz and Holt (1995): rescue is either caused by minute changes in allele frequencies across many loci in sexuals (i.e., the infinitesimal model; Fisher 1918) or by the substitution of a single large effect ‘resistance’ mutation (e.g., one locus, two allele models). We therefore largely lack a theoretical framework for the genetic basis of evolutionary rescue that captures the arguably more realistic situation where an intermediate number of mutations are at play (but see exceptions below). The near absence of such a framework prevents us from predicting the number of mutations that evolutionary rescue will take and the distribution of their effect sizes. The existence of a more complete framework could therefore provide valuable information for those investigating the genetic basis of drug resistance (e.g., the expected number and effect sizes of mutations) and would extend our understanding of the genetic basis of adaptation to cases of non-equilibrial demography (i.e., rapid evolution and “eco-evo” dynamics).

Despite these gaps in the theory on the genetic basis of evolutionary rescue, there is a wealth of data. For example, the genetic basis of resistance to a variety of drugs is known in many species of bacteria (reviewed in MacLean *et al.* 2010), fungi (reviewed in Robbins *et al.* 2017), and viruses (reviewed in *Yilmaz et al.* 2016). This abundance of data reflects both the applied need to prevent drug resistance and the relative ease of isolating the genotypes that survive (hereafter “rescue genotypes”), e.g., in a Luria-Delbrück fluctuation assay (reviewed in Bataillon and Bailey 2014). Assaying fitness in the environment used to isolate mutants (e.g., in the drug) then provides the distribution of fitness effects of potential rescue genotypes. Additional data on the genetic basis of drug resistance arise from the construction of mutant libraries (e.g., Weinreich *et al.* 2006) and the sequencing of natural populations (e.g., Pennings *et al.* 2014). Together, the data show that resistance often appears to arise by a single mutation (e.g., MacLean and Buckling 2009; Lindsey *et al.* 2013; Gerstein *et al.* 2012) but not always (e.g., Bataillon *et al.* 2011; Pennings *et al.* 2014; Gerstein *et al.* 2015; Williams and Pennings 2019). The data also indicate that the fitness effect of rescue genotypes is more often large than small, creating a hump-shaped distribution of selection coefficients (e.g., Kassen and Bataillon 2006; MacLean and Buckling 2009; Gerstein *et al.* 2012; Lindsey *et al.* 2013; Gerstein *et al.* 2015) that is similar in shape to that proposed by Kimura (1983) (see Orr 1998, for more discussion) but with a lower bound that is often much greater than zero.

Theory on evolutionary rescue (reviewed in *Alexander et al.* 2014) has primarily focused on the probability of rescue rather than its genetic basis. However, a few studies have varied the potential genetic basis enough to make some inference about how evolutionary rescue is likely to happen. For instance, in the context of pathogen host-switching, Antia *et al.* (2003) numerically explored the probability of rescue starting from a single ancestral individual when *k* sequential mutations are required for a positive growth rate, each mutation occurring from the previous genotype with the same probability and all intermediate genotypes being selectively neutral. The authors found that rescue became less likely as the number of intermediate mutations increased, suggesting that rescue will generally proceed by the fewest possible mutations. This framework was expanded greatly by Iwasa *et al.* (2004a), who allowed for arbitrary mutational networks (i.e., different mutation rates between any two genotypes) and standing genetic variation in the ancestral population. Assuming the probability of mutation between any two genotypes is of the same order, they showed that genetic paths with fewer mutational steps contributed more to the total probability of rescue, again suggesting rescue will occur by the fewest possible mutations. Iwasa *et al.* (2004a) also found that multiple simultaneous mutations (i.e., arising in the same meiosis) can contribute more to rescue than paths that gain these same mutations sequentially (i.e., over multiple generations) when the growth rates of the intermediate mutations are small enough, suggesting that rare large mutations can be the most likely path to rescue when the population is very maladapted or there is a fitness valley separating the wildtype and rescue genotype. This point was also demonstrated by Alexander and Day (2010), who emphasized that multiple simultaneous mutations become the dominant path to rescue in the most challenging environments. As a counterpoint, Uecker and Hermisson (2016) explored a greater range of fitness values in a two-locus two-allele model, showing that, with standing genetic variation, rescue by sequential mutations at two loci (two mutational steps) can be more likely than rescue by mutation at a single locus (one simultaneous mutational step), particularly when the wildtype is very maladapted, where the single mutants can act as a buffer in the face of environmental change. In summary, current theory indicates that the genetic basis of rescue hinges on the chosen set of genotypes, their fitnesses, and the mutation rates between them. So far these choices have been in large part arbitrary or chosen for mathematical convenience.

Here we follow the lead of Anciaux *et al.* (2018) in allowing the genotypes that contribute to rescue, as well as their fitnesses and the mutational distribution, to arise from an empirically-justified fitness-landscape model (Tenaillon 2014). In particular, we use Fisher’s geometric model to describe adaptation following an abrupt environmental change that instigates population decline. There are two key differences between this approach and earlier models using Fisher’s geometric model (e.g., Orr 1998): here 1) the dynamics of each genotype depends on their absolute fitness (instead of only on their relative fitness) and 2) multiple mutations can segregate simultaneously (instead of assuming only sequential fixation), allowing multiple mutations to fix – and in our case, rescue the population – together as a single haplotype (i.e., stochastic tunnelling, Iwasa *et al.* 2004b). In this non-equilibrium scenario, variation in absolute fitness, which allows population size to vary, can create feedbacks between demography and evolution, which could strongly impact the genetic basis of adaptation relative to the constant population size case. In contrast to Anciaux *et al.* (2018), our focus here is on the genetic basis of evolutionary rescue and we also explore the possibility of rescue by mutant haplotypes containing more than one mutation. In particular, we ask: (1) How many mutational steps is evolutionary rescue likely to take? and (2) What is the expected distribution of fitness effects of the surviving genotypes and their component mutations?

We first introduce the modelling framework before summarizing our main results. We then present the mathematical analyses we have used to understand these results and end with a discussion of our key findings.

## Data availability

Code used to derive analytical and numerical results and produce figures (referred to here as File S1; *Mathematica*, version 9.0; Wolfram Research Inc. 2012) and code used to create individual-based simulation data (Python, version 3.5; Python Software Foundation), as well as simulation data and freely accessible versions of File S1 (CDF and PDF), are available at https://github.com/mmosmond/GeneticBasisOfRescue.

## Model

### Fisher’s geometric model

We map genotype to phenotype to fitness using Fisher’s geometric model, originally introduced by Fisher (1930, p. 38-41) and reviewed by Tenaillon (2014). In this model each genotype is characterized by a point in *n*-dimensional phenotypic space, 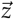. We ignore environmental effects, and thus the phenotype is the breeding value. At any given time there is a phenotype, 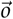, that has maximum fitness and fitness declines monotonically as phenotypes depart from 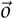. We assume that *n* phenotypic axes can be chosen and scaled such that fitness is described by a multivariate Gaussian function with variance 1 in each dimension, no covariance, and height *W*_*max*_ (which can always be done when considering genotypes close enough to an non-degenerate optimum; Martin 2014). Thus the fitness of phenotype 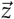 is 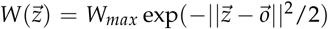, where 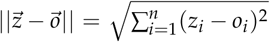 is the Euclidean distance of 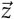 from the optimum, 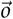. Here we are interested in absolute fitness; we take 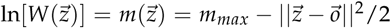 to be the continuous-time growth rate (*m* is for *M*althusian fitness) of phenotype 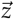. We ignore density- and frequency-dependence in 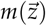 for simplicity. The fitness effect, i.e., selection coefficient, of phenotype *z′* relative to *z* in a continuous-time model is exactly *s* = log[*W*(*z′*)/*W*(*z*)] = *m*(*z*′) − *m*(*z*) (Martin and Lenormand 2015). This is approximately equal to the selection coefficient in discrete time (*W*(*z′*)/*W*(*z*) − 1) when selection is weak (*W*(*z′*) − *W*(*z*) << 1).

To make analytical progress we use the isotropic version of Fisher’s geometric model, where mutations (in addition to selection) are assumed to be uncorrelated across the scaled traits. Universal pleiotropy is also assumed, so that each mutation affects all scaled phenotypes. In particular we use the “classic” form of Fisher’s geometric model (Harmand *et al.* 2017), where the probability density function of a mutant phenotype is multivariate normal, centred on the current phenotype, with variance *λ* in each dimension and no covariance. Using a probability density function of mutant phenotypes implies a continuum-of-alleles (Kimura 1965), i.e., phenotype is continuous and each mutation is unique. Mutations are assumed to be additive in phenotype, which induces epistasis in fitness (as well as dominance under diploid selection), as fitness is a non-linear function of phenotype. We assume asexual reproduction, i.e., no recombination, which is appropriate for many cases of antimicrobial drug resistance and experimental evolution, while recognizing the value of expanding this work to sexual populations.

An obvious and important extension would be to relax the simplifying assumptions of isotropy and universal pleiotropy, which we leave for future work. Note that mild anisotropy yields the same bulk distribution of fitness effects as an isotropic model with fewer dimensions (Martin and Lenormand 2006), but this does not extend to the tails of the distribution. Therefore, whether anisotropy can be reduced to isotropy with fewer dimensions in the case of evolutionary rescue, where the tails are essential, is unknown. In the Discussion we briefly explore the effects of non-Gaussian distributions of mutant phenotypes.

Given this phenotype-to-fitness mapping and phenotypic distribution of new mutations, the distribution of fitness effects (and therefore growth rates) of new mutations can be derived exactly. Let *m* be the growth rate of some particular focal genotype and *m*′ the growth rate of a mutant immediately derived from it. Then let *s*_*o*_ = *m*_*max*_−*m* be the selective effect of a mutant with the optimum genotype and *s* = *m*′ − *m* the selective effect of the mutant with growth rate *m*′. The probability density function of the selective effects of new mutations, *s*, is then given by equation 3 in Martin and Lenormand (2015). Converting fitness effects to growth rate (*m*′ = *s* + *m*), the probability density function for mutant growth rate *m*′ from an ancestor with growth rate *m* is (cf. equation 2 in Anciaux *et al.* 2018)

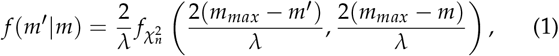

where 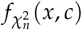 is the probability density function over positive real numbers *x* of 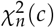, a non-central chi-square deviate with *n* degrees of freedom and noncentrality *c* > 0 (equation 26.4.25 in Abramowitz and Stegun 1972).

### Lifecycle

We are envisioning a scenario where *N*_0_ wildtype individuals, each of which have phenotype 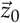, experience an environmental change, causing population decline, 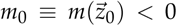. Each generation, an individual with phenotype 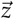 produces a Poisson number of offspring, with mean 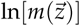, and dies. This process implicitly assumes no interaction between individuals, i.e., a branching process with density- and frequency-independent growth and fitness and no clonal interference. Each offspring mutates with probability *U* (we ignore the possibility of multiple simultaneous mutations within a single genome), and mutations are distributed as described above (see Fisher’s geometric model).

### Simulation procedure

We ran individual-based simulations of the above process to compare with our numeric and analytic results. Populations were considered rescued when there were ≥ 1000 individuals (Figures 1–3) or ≥ 100 individuals (Figures 6–7, S1, and S3) with positive growth rates (all other replicates went extinct). The most common genotype at the time of rescue was considered the rescue genotype, and the number of mutational steps to rescue was set as the number of mutations in that genotype.

**Figure 1.**
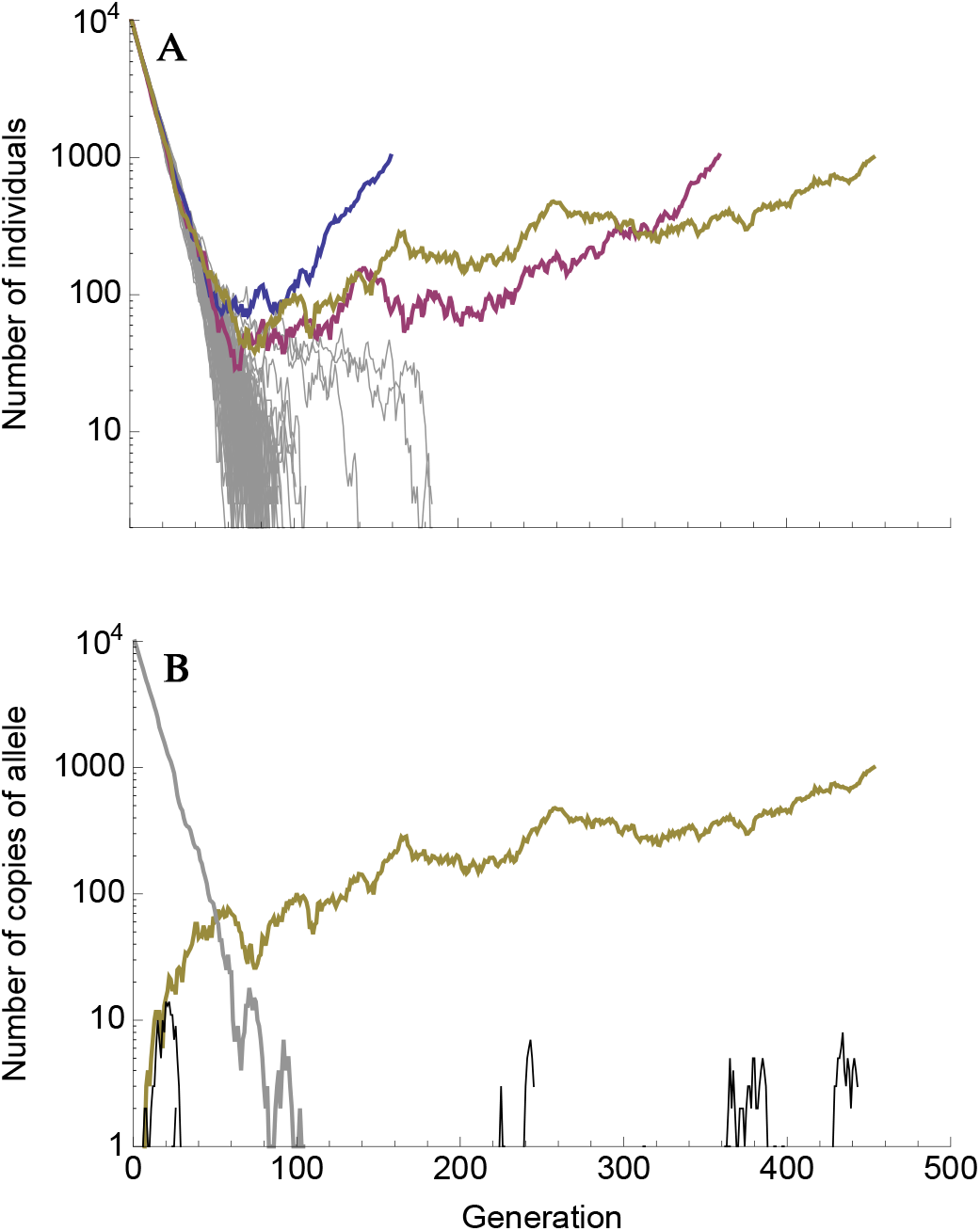
Typical dynamics with a relatively slow wildtype decline and a small mutation rate (*m*_0_ = −0.1, *U* = 10^−4^). **(A)** Population size trajectories on a log scale. Each line is a unique replicate simulation (100 replicates). Replicates that went extinct are grey, replicates that were rescued are in colour (and are roughly V-shaped). **(B)** The number of individuals with a given derived allele, again on a log scale, for the yellow replicate in **A**. The number of individuals without any derived alleles (wildtypes) is shown in grey, the rescue mutation is shown in yellow, and all other mutations are shown in black. Other parameters: *n* = 4, *λ* = 0.005, *m*_*max*_ = 0.5.

### Probability of rescue

Let *p*_0_ be the probability that a given wildtype individual is “successful”, i.e., has descendants that rescue the population. The probability of rescue is then one minus the probability that none of the initial wildtype individuals are successful,

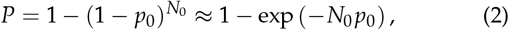

where the approximation assumes small *p*_0_ and large *N*_0_. What remains is to find *p*_0_.

## Summary of Results

We start with a heuristic explanation of our main results before turning to more detailed derivations in the next section.

### Rescue by multiple mutations

A characteristic pattern of evolutionary rescue is a “U”-shaped population size trajectory (e.g., Orr and Unckless 2014). This is the result of an exponentially-declining wildtype genotype being replaced by an exponentially-increasing mutant genotype. On a log scale this population size trajectory becomes “V”-shaped (we denote it a ‘V-shaped log-trajectory’). On this scale, the population declines at a constant rate (producing a line with slope *m*_0_ < 0) until the growing mutant subpopulation becomes relatively common, at which point the population begins growing at a constant rate (a line with slope *m*_1_ > 0). This characteristic V-shaped log-trajectory is observed in many of our simulations where evolutionary rescue occurs (Figure 1A). Alternatively, when the wildtype declines faster and the mutation rate is larger we sometimes see ‘U-shaped log-trajectories’ (e.g., the red and blue replicates in Figure 2A). Here there are three phases instead of two; the initial rate of decline (a line with slope *m*_0_ < 0) is first reduced (transitioning to a line with slope *m*_1_ < 0) before the population begins growing (a line with slope *m*_2_ > 0).

**Figure 2.**
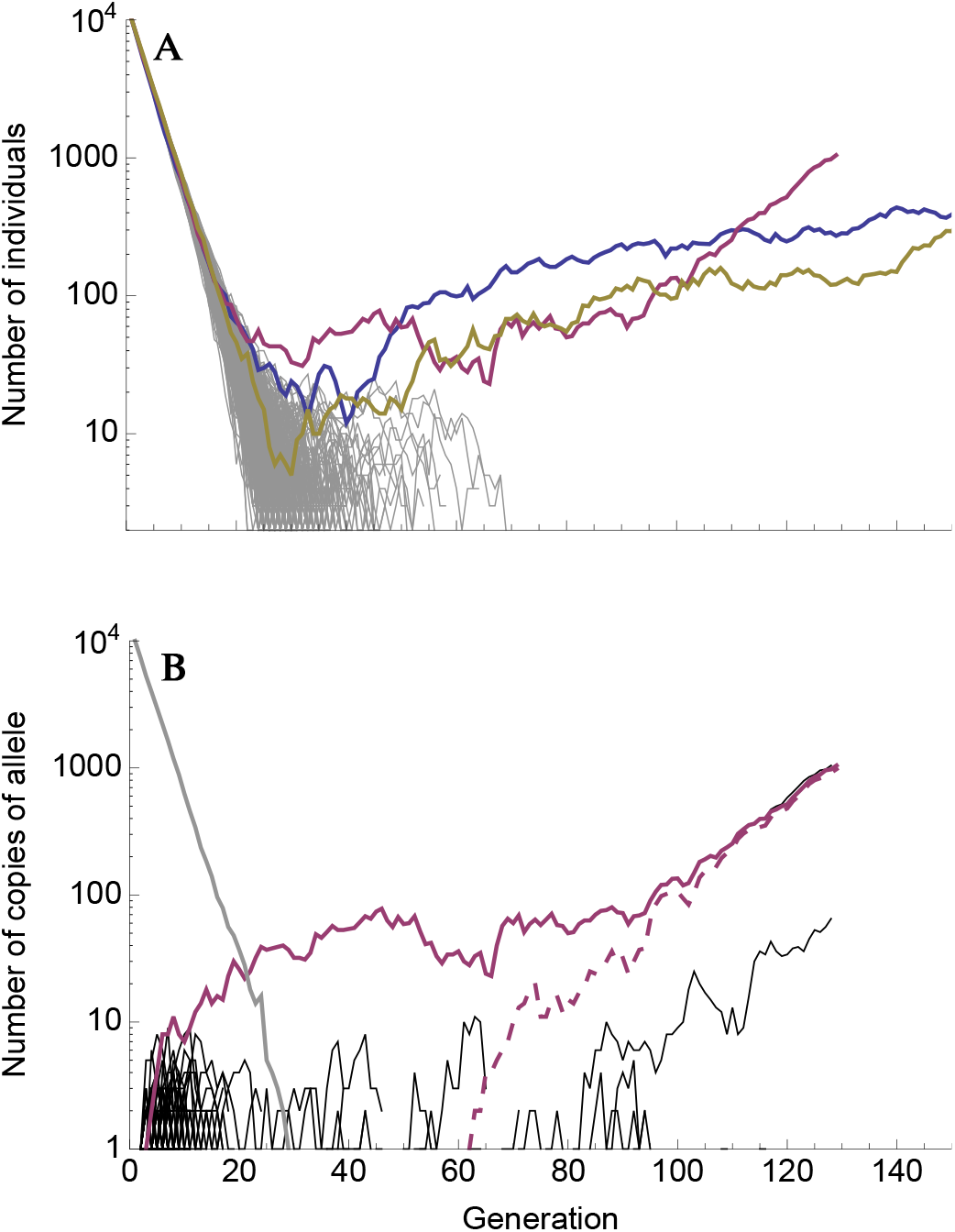
Typical dynamics with a relatively fast wildtype decline and a large mutation rate (*m*_0_ = 0.3, *U* = 10^−2^). **(A)** Population size trajectories on a log scale. Each line is a unique replicate simulation (500 replicates). Replicates that went extinct are grey, replicates that were rescued are in colour. Note that the blue and red replicates are cases of 2-step rescue (and roughly U-shaped), while the yellow replicate is 1-step rescue (and therefore V-shaped). **(B)** The number of individuals with a given derived allele, again on a log scale, for the red replicate in **A**. The number of individuals without any derived alleles (wildtypes) is shown in grey, the rescue mutations are shown in red, and all other mutations in black. Here a single mutant with growth rate less than zero arises early and outlives the wildtype (solid red). A second mutation then arises on that background (dashed red), making a double mutant with a growth rate greater than zero that rescues the population. Other parameters: *n* = 4, *λ* = 0.005, *m*_*max*_ = 0.5.

As expected, V-shaped log-trajectories are the result of a single mutation creating a genotype with a positive growth rate that escapes loss when rare and rescues the population (Figure 1B), i.e., 1-step rescue. U-shaped log-trajectories, on the other hand, occur when a single mutation creates a genotype with a negative (or potentially very small positive) growth rate, itself doomed to extinction, which out-persists the wildtype and gives rise to a double mutant genotype that rescues the population (Figure 2B), i.e., 2-step rescue. These two types of rescue comprise the overwhelming majority of rescue events observed in our simulations, across a wide range of wildtype decline rates (e.g., Figure 3).

**Figure 3.**
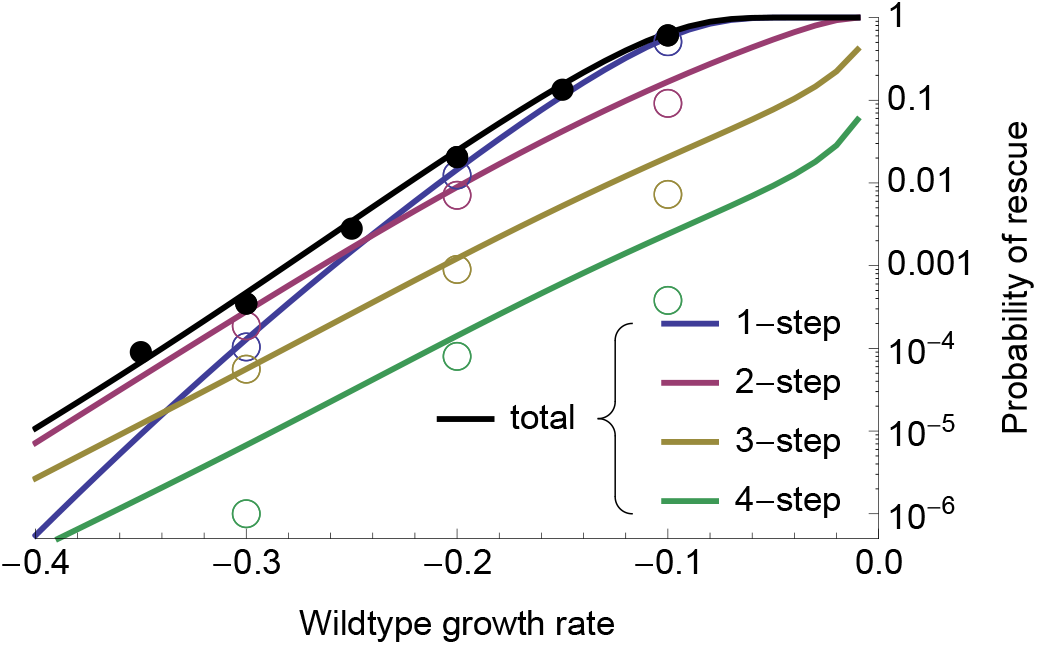
The probability of evolutionary rescue as a function of initial maladaptation. Shown are the probabilities of 1-, 2-, 3-, and 4-step rescue (using Equations 2–7), as well as the probability of rescue by up to 4 mutational steps (“total”, using 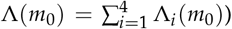. Circles are individual-based simulation results (ranging from 10^5^ to 10^6^ replicates per wildtype growth rate). Open circles denote the fraction of simulations where the rescue genotype (see Simulation procedure) had a given number of mutations and closed circles are the sum of these fractions. Parameters: *N*_0_ = 10^4^, *U* = 2 × 10^−3^, *n* = 4, *λ* = 0.005, *m*_*max*_ = 0.5.

In the text, we focus on low to moderate mutation rates affecting growth rate. With sufficiently high mutation rates rescue by 3 or more mutations comes to dominate (Figure S1). It has recently been suggested that when the mutation rate, *U*, is substantially less than a critical value, *U*_*C*_ = *λn*^2^/4, we are in a “strong selection, weak mutation” regime where selection is strong enough relative to mutation that essentially all mutations arise on a wildtype background (Martin and Roques 2016), consistent with the House of Cards approximation (Turelli 1984, 1985). Thus in this regime rescue tends to occur by a single mutation of large effect (Anciaux *et al.* 2018). In the other extreme, when *U* >> *U*_*C*_, we are in a “weak selection, strong mutation” regime where selection is weak enough relative to mutation that many cosegregating mutations are present within each genome, creating a multivariate normal phenotypic distribution (Martin and Roques 2016), consistent with the Gaussian approximation (Kimura 1965; Lande 1980). Thus in this regime rescue tends to occur by many mutations of small effect (Anciaux *et al.* 2019). As shown in Figure 3 (where *U* = *U*_*C*_ /10) and Figure S1 (where *U*_*C*_ = 0.02), rescue by a small number of mutations (but more than one) can become commonplace in the transition zone (where *U* is neither much smaller or much larger than *U*_*C*_), where there are often a considerable number of cosegregating mutations (e.g., Figure 2B, where *U* = *U*_*C*_/2).

### The probability of k-step rescue

Approximations for the probability of 1-step rescue under the strong selection, weak mutation regime were derived by Anciaux *et al.* (2018). Here we extend this study by exploring the contribution of *k*-step rescue, deriving approximations for the probability of such events, as well as dissecting the genetic basis of both 1- and 2-step rescue in terms of the distribution of fitness effects of rescue genotypes and their component mutations.

Although requiring a sufficiently beneficial mutation to arise on a rare mutant genotype doomed to extinction, multi-step evolutionary rescue can be the dominant form of rescue when the wildtype is sufficiently maladapted (Figures 3 and S1). Indeed, on this fitness landscape, the probability of producing a rescue genotype in one mutational step mutant drops very sharply with maladaptation (Anciaux *et al.* 2018); the probability of multi-step rescue declines more slowly as mutants with intermediate growth rates can be a “springboard” – albeit not always a very bouncy one – from which rescue mutants are produced. These intermediates contribute more as mutation rates and the decline rate of the wildtype increase (Figures 3 and S1), the former because double mutants become more likely and the latter because the springboard becomes more necessary. With a large enough number of wildtype individuals or a high enough mutation rate (Figure S1), multi-step rescue can not only be more likely than 1-step, but also very likely in an absolute sense.

### Classifying 2-step rescue regimes

2-step rescue can occur through first-step mutants with a wide range of growth rates. As shown below (see Approximating the probability of 2-step rescue), these first-step mutants can be divided into three regimes: “sufficiently subcritical”, “sufficiently critical”, and “sufficiently supercritical” (we will often drop “sufficiently” for brevity; Figure 4). Sufficiently critical first-step mutants are defined by having growth rates close enough to zero that the most likely way for such a mutation to lead to 2-step rescue is for it to persist for such an unusually long period of time, and accordingly grow to such an unusually large sub-population size, that it will almost certainly produce successful double mutants. Sufficiently subcritical first-step mutants are then defined by having growth rates that are negative enough to almost certainly prevent such long persistence times. Instead, these mutations tend to persist for an expected number of generations, proportional to the inverse of their growth rate (1/|*m*|), while maintaining relatively small subpopulation sizes (on the order of one individual per generation). Mutations conferring a positive growth rate can also go extinct, and thus can also act as springboards to rescue. Conditioned on extinction, supercritical mutations behave like subcritical mutations with a growth rate of the same absolute value (Maruyama and Kimura 1974). Sufficiently supercritical first-step mutants are therefore defined analogously to subcritical first-step mutants, having positive (rather than negative) growth rates that are large enough to prevent sufficiently long persistence times once conditioned on extinction. Despite having similar extinction trajectories as subcritical mutations, ‘doomed’ supercritical mutations arise less frequently by mutation from the wildtype but mutate to rescue genotypes at a higher rate. Overall, they too can contribute substantially to rescue. Note that supercritical 2-step rescue is not 1-step rescue with subsequent adaptation as we condition on the first-step mutation going extinct in the absence of the second mutation. However, empirically it will be impossible to tell if the first-step mutation was indeed doomed to extinction if it is found to have a positive growth rate in the selective environment.

**Figure 4.**
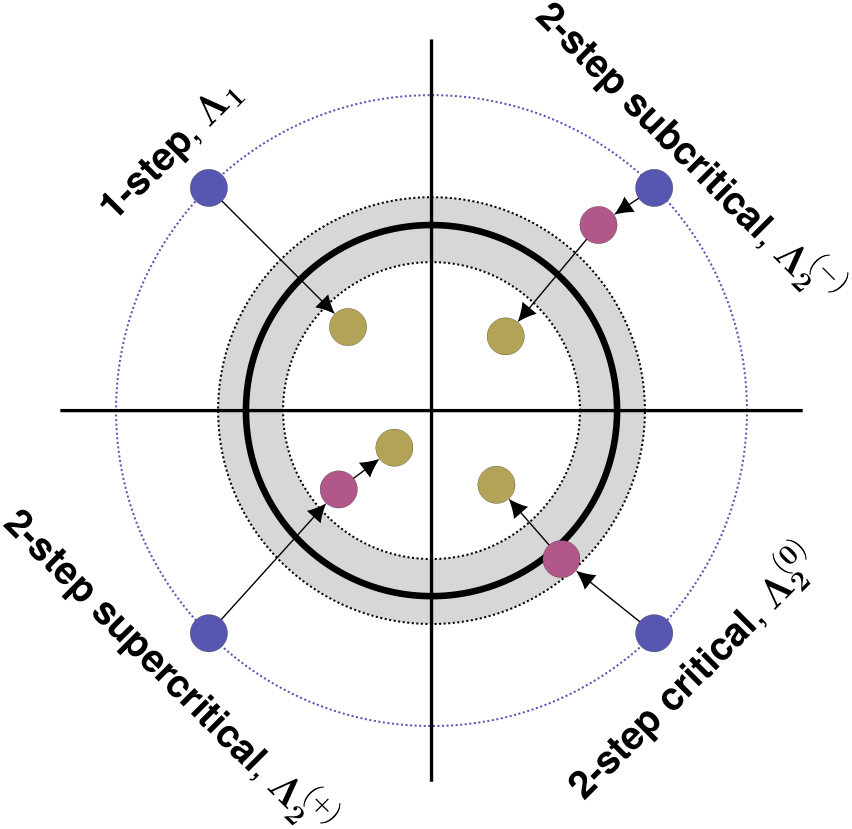
1- and 2-step genetic paths to evolutionary rescue. Here we show an *n* = 2 dimensional phenotypic landscape. Continuous-time (Malthusian) growth rate (*m*) declines quadratically from the centre, becoming negative outside the thick black line. The grey zone indicates where growth rates are “sufficiently critical” (see text for details). Blue circles show wildtype phenotypes, red circles show intermediate first-step mutations, and yellow circles show the phenotypes of rescue genotypes.

The relative contribution of each regime changes with both the initial degree of maladaptation and the mutation rate (Figures 5 and S2). When the wildtype is very maladapted (relative to mutational variance), most 2-step rescue events occur through subcritical first-step mutants (Figure 5A), which arise at a higher rate than critical or supercritical mutants and yet persist longer than the wildtype. When the wildtype is less maladapted, however, critical and supercritical mutations become increasingly likely to arise and contribute to 2-step rescue, both due to their closer proximity to the wildtype in phenotypic space as well as the slower decline of the wildtype increasing the cumulative number of mutations that occur. The mutation rate also plays an interesting role in determining the relative contributions of each regime (Figures 5B and S2). When mutations are rare, only first-step mutations that are very nearly neutral (*m* ~ 0) will persist long enough to give rise to a 2-step rescue mutation. As the mutation rate increases, however, the range of first-step mutant growth rates that can persist long enough to lead to 2-step rescue widens because fewer individuals carrying the first-step mutation are needed before a successful double mutant arises.

**Figure 5.**
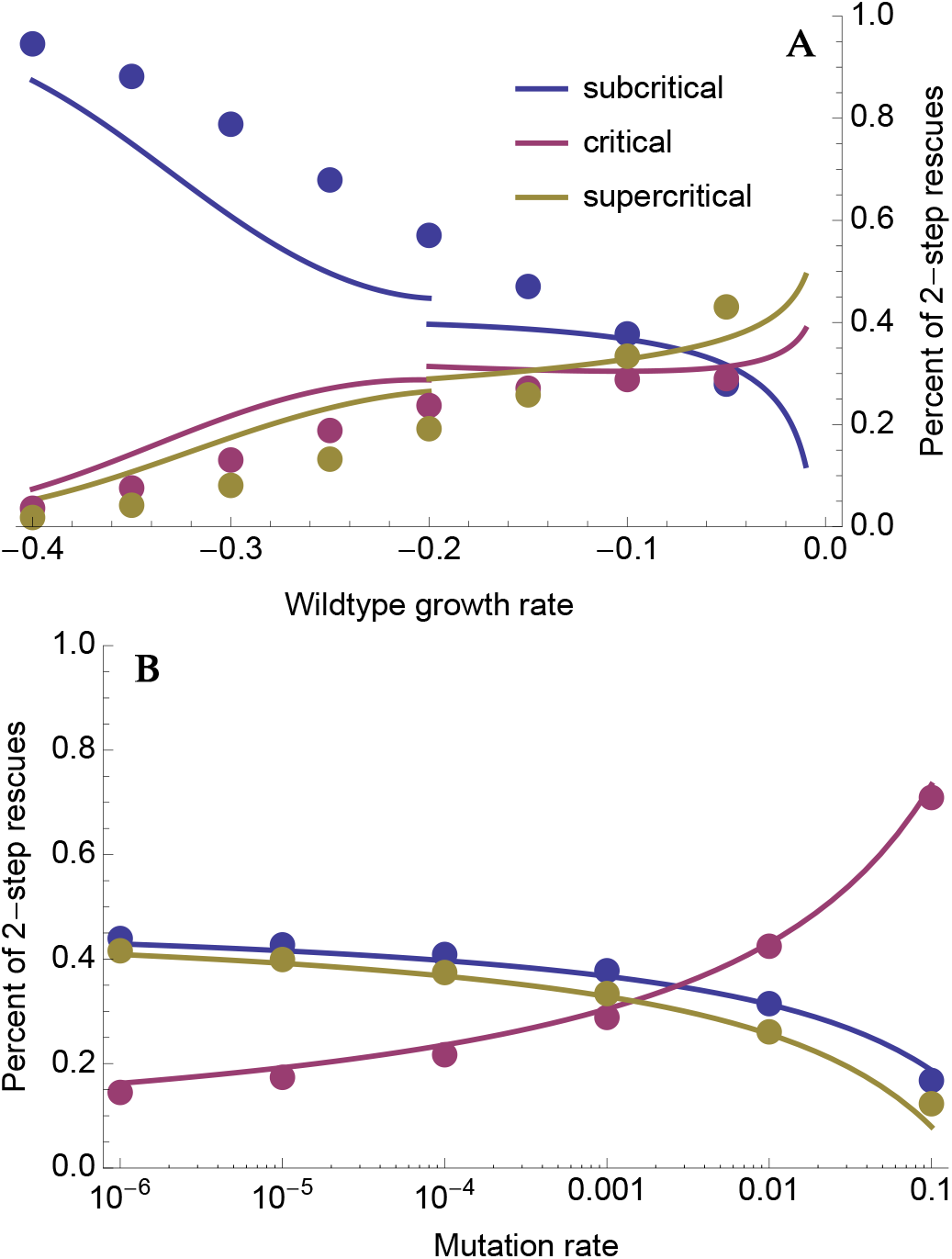
The relative contribution of sufficiently subcritical, critical, and supercritical single mutants to 2-step rescue. The curves are drawn using Equations 10–14 (Equation 12 is used for *m*_0_ < 0.2 while Equation 13 is used for *m*_0_ > 0.2). The dots are numerical evaluations of Equation 8. Parameters: *n* = 4, *λ* = 0.005, *m*_*max*_ = 0.5, (**A**) *U* = 10^−3^, (**B**) *m*_0_ = −0.1.

### The distribution of fitness effects among rescue mutations

Mutants causing 1-step rescue have growth rates that cluster around small positive values (*m* ≳ 0; blue curves in Figure 6). Consequently, the distribution of fitness effects (DFE) among these rescue mutants is shifted to the right relative to mutations that establish in a population of constant size (compare solid blue and gray curves in Figure 6), with a DFE beginning at *s* = *m* − *m*_0_ ≥ −*m*_0_ > 0 rather than *s* = 0 (Kimura 1983). As a result of this increased threshold, the 1-step rescue DFE has a smaller variance than both the DFE of random mutations and the DFE of mutations that establish in a constant population (compare blue and gray curves in Figure 6). Further, while the variance in the DFE of random mutations and of those that establish in a population of constant size increases slightly with initial maladaptation (due to the curvature of the phenotype-to-fitness function), the variance in the 1-step rescue DFE decreases substantially (compare panels in Figure 6), as rescue becomes restricted to a rapidly decreasing proportion of the available mutants.

**Figure 6.**
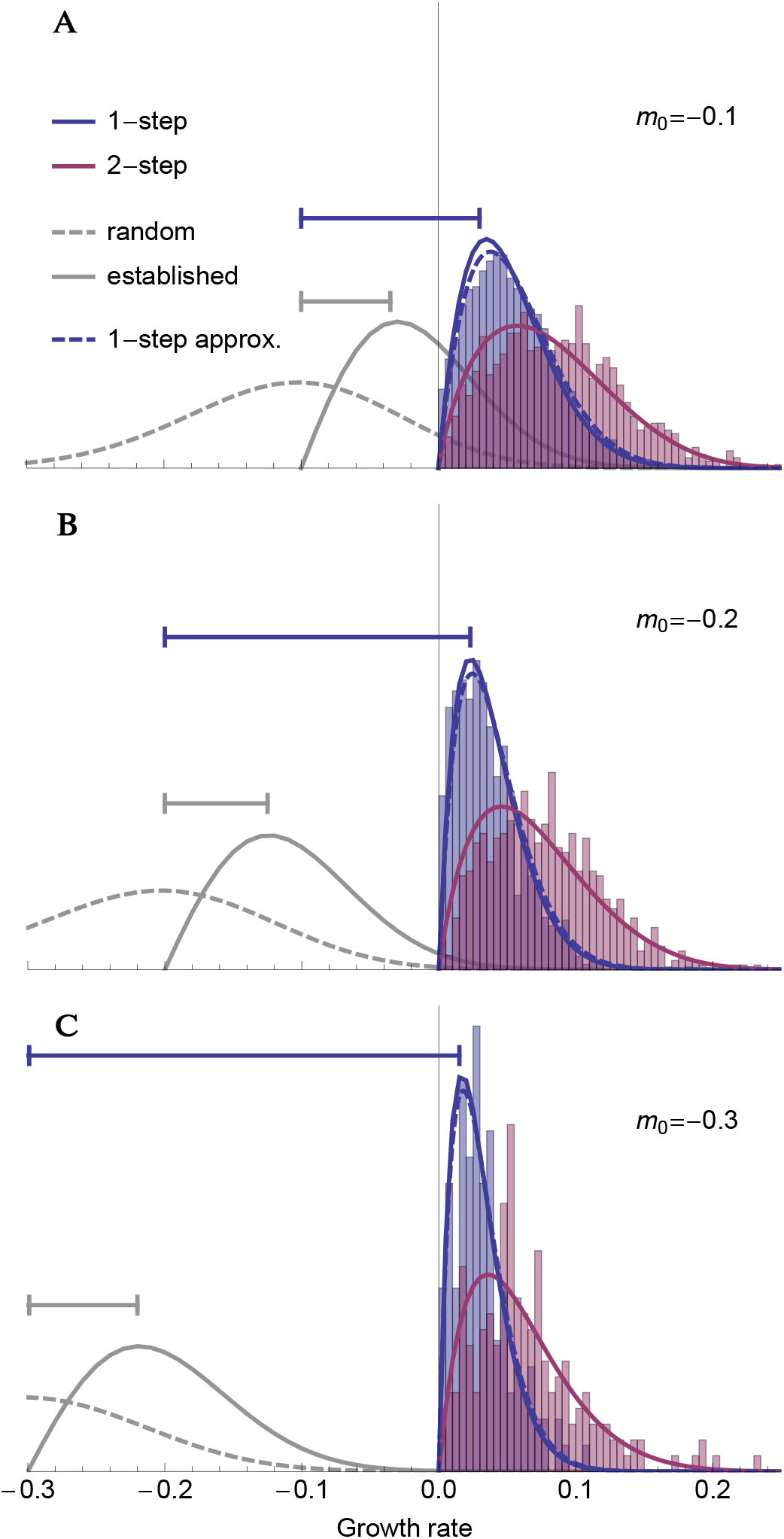
The distribution of growth rates among rescue genotypes under 1-step (blue; Equation 15 solid and 16 dashed) and 2-step (red; Equation 17) rescue for three different levels of initial maladaptation. For comparison, the distribution of random mutations (dashed; Equation 1) and the distribution of beneficial mutations that establish in a population of constant size (solid grey; Equation 1 times Equation 4 and normalized) are shown. Intervals (horizontal lines) indicate the size of the most common fitness effect (*s* = *m*_0_ − *m*) in a population of constant size (grey) and in 1-step rescue (blue). The histograms show the distribution of growth rates among rescue genotypes observed across (**A**) 10^4^, (**B**) 10^5^, and (**C**) 10^6^ simulated replicates. Other parameters: *N*_0_ = 10^4^, *U* = 2 × 10^−3^, *n* = 4, *λ* = 0.005, *m*_*max*_ = 0.5.

The DFE of genotypes that cause 2-step rescue (the combined effect of two mutations) is also clustered at small positive growth rates, but it has a variance that is less affected by the rate of wild-type decline (red curves in Figure 6). This is because double mutant rescue genotypes are created via first-step mutant genotypes that have larger growth rates than the wildtype (i.e., are closer to the optimum), allowing them to create double mutants with a larger range of positive growth rates.

Finally, we can also look at the distribution of growth rates among first-step mutations that lead to 2-step rescue, i.e., ‘springboard mutants’ (Figures 7 and S2). Here there are two main factors to consider: 1) the probability that a mutation with a given growth rate arises on the wildtype background but does not by itself rescue the population and 2) the probability that such a mutation persists long enough for a sufficiently beneficial second mutation to arise on that same background and together rescue the population. Subcritical mutations conferring growth rates closer to zero persist longer but are less likely to arise from the wildtype, creating a trade-off between mutational input and the probability of rescue that can lead to a wide distribution of contributing subcritical growth rates (blue shading in Figure 7). In contrast, supercritical mutations with growth rates nearer to zero are more likely arise by mutation, to go extinct in the absence of further mutation, and to persist for longer once conditioned on extinction, together creating a relatively narrow distribution of contributing supercritical growth rates (yellow shading in Figure 7). As explained above, increasing the rate of wildtype decline (or decreasing the rate of mutation) increases the contribution of subcritical first-step mutants and the importance of mutational input, lowering the mode and increasing the variance of the first-step DFE (compare panels in Figure 7).

**Figure 7.**
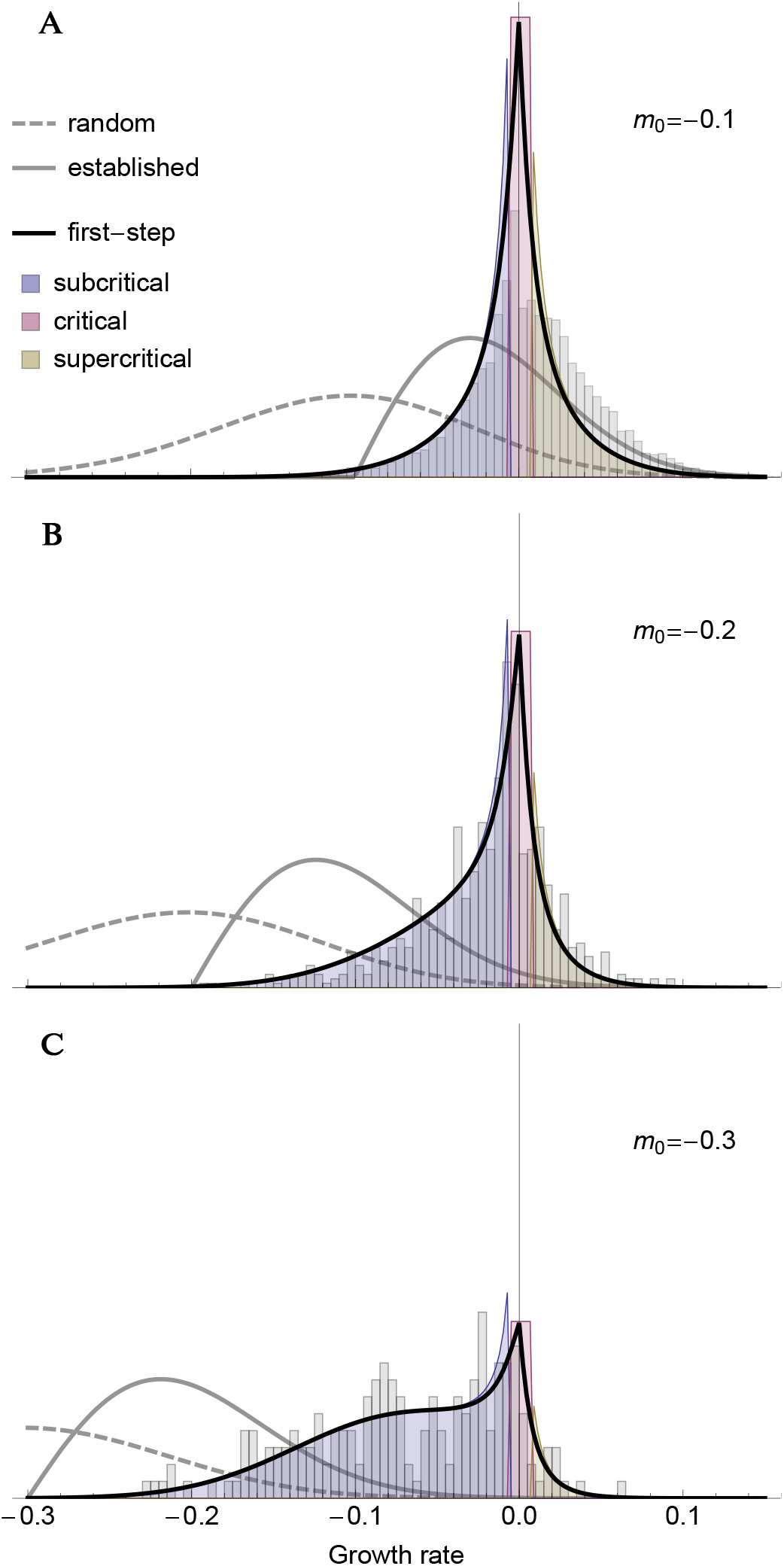
The distribution of growth rates among first-step mutations that lead to 2-step rescue (black; Equation 18) for three different levels of initial maladaptation. Shading represents our sufficiently subcritical approximation (blue; replacing *p*(*m*, Λ_1_(*m*)) with Λ_1_(*m*)/|*m*| in the numerator of Equation 18), our sufficiently critical approximation (red; using 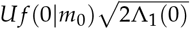 as the numerator in Equation 18), and our sufficiently supercritical approximation (yellow; replacing *p*(*m*, Λ_1_(*m*)) with Λ_1_(*m*)/|*m*| in the numerator of Equation 18). The histograms show the distribution of growth rates among first-step mutations in rescue genotypes with 2 mutations observed across (**A**, **B**) 10^5^ or (**C**) 10^6^ simulated replicates. We hypothesize that the overabundance of supercriticals (especially in panel **A**) is likely due to us sampling only the most common rescue genotype in each replicate, which is not necessarily the first genotype that rescues. See Figure 6 for additional details.

Note that, given 2-step rescue, the growth rate of both the first-step and second-step mutation may be negative when considered by themselves in the wildtype background. This potentially obscures empirical detection of the individual mutations involved in evolutionary rescue.

## Mathematical Analysis

### The probability of k-step rescue

Generic expressions for the probability of 1- and 2-step rescue were given by Martin *et al.* (2013), using a diffusion approximation of the underlying demographics. The key result that we will use is the probability that a single copy of a genotype with growth rate *m*, itself fated for extinction but which produces rescue mutants at rate Λ(*m*), rescues the population (equation S1.5 in Martin *et al.* 2013). With our lifecycle this is (c.f., equation A.3 in Iwasa *et al.* 2004a)

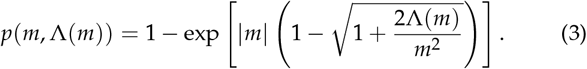

We can therefore use *p*_0_ = *p*(*m*_0_, Λ(*m*_0_)) as the probability that a wildtype individual has descendants that rescue the population and what remains in calculating the total probability of rescue (Equation 2) is Λ(*m*_0_). We break this down by letting Λ_*i*_ (*m*) be the rate at which rescue genotypes with *i* mutations are created; the total probability of rescue is then given by Equation 2 with 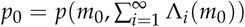.

In 1-step rescue, Λ_1_(*m*_0_) is just the rate of production of rescue mutants directly from a wildtype genotype. This is the probability that a wildtype gives rise to a mutant with growth rate *m* (given by *Uf* (*m*|*m*_0_)) times the probability that a genotype with growth rate *m* establishes. Again approximating our discrete time process with a diffusion process, the probability that a lineage with growth rate *m* << 1 establishes, ignoring further mutation, is (e.g., Martin *et al.* 2013)

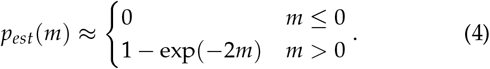

This reduces to the 2(*s* + *m*_0_) result in Otto and Whitlock (1997) when *m* = *s* + *m*_0_ is small, which further reduces to 2*s* in a population of constant size, where *m*_0_ = 0 (Haldane 1927). Using this, the rate of 1-step rescue is

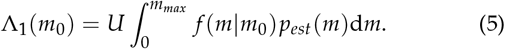

**Table 1.**
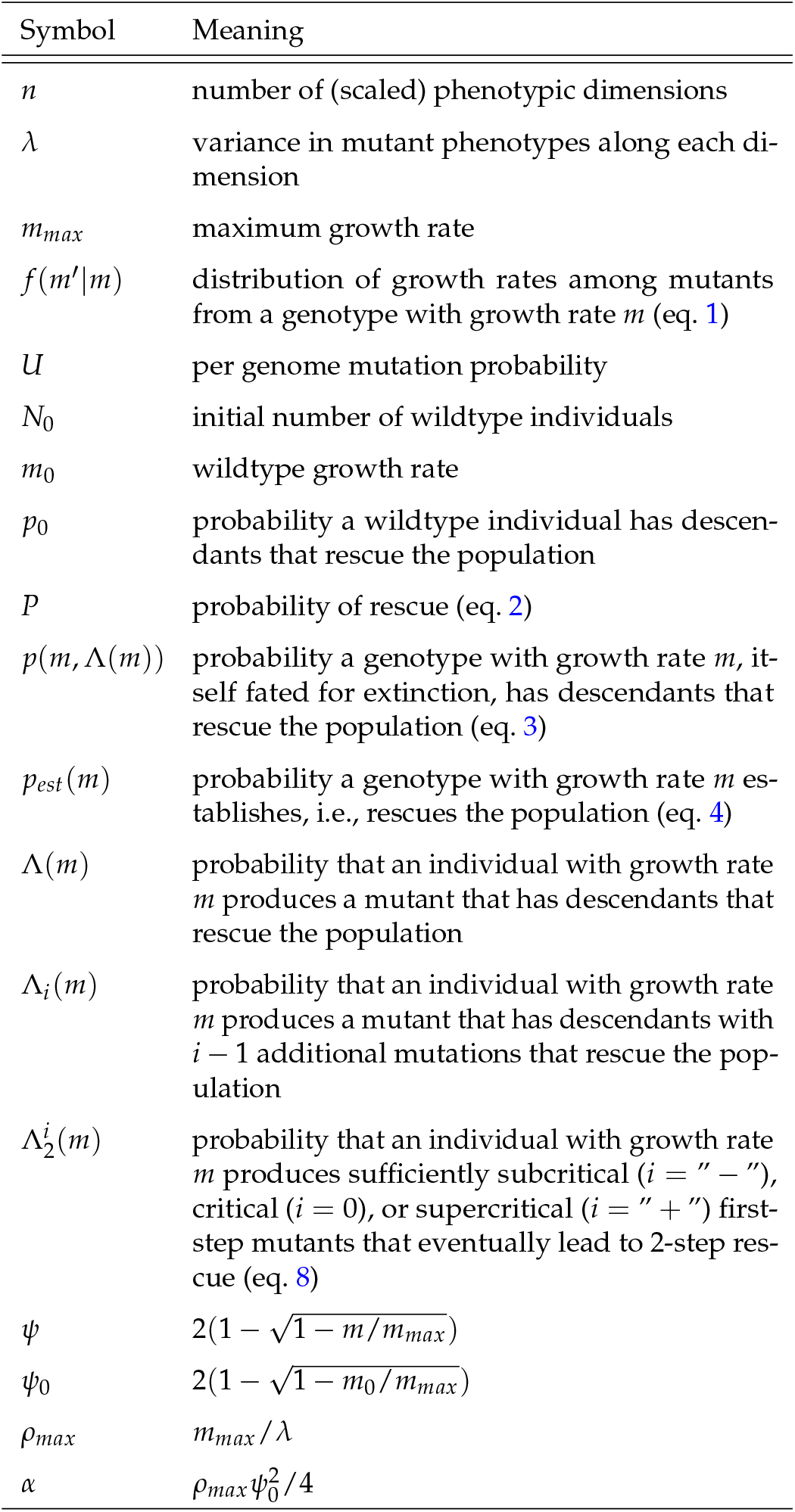
Frequently used notation.

Taking the first order approximation of *p*(*m*_0_, Λ_1_(*m*_0_)) with 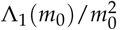 small gives the probability of 1-step rescue (equation 5 of Anciaux *et al.* 2018), which effectively assumes deterministic wildtype decline. For completeness we rederive their closed-form approximation in File S1 (and give the results in the Appendix, see Approximating the probability of 1-step rescue).

The probability of 2-step rescue is only slightly more complicated. Here Λ_2_(*m*_0_) is the probability that a mutation arising on the wildtype background creates a genotype that is also fated for extinction but persists long enough for a second mutation to arise on this mutant background, creating a double mutant genotype that rescues the population. We therefore have

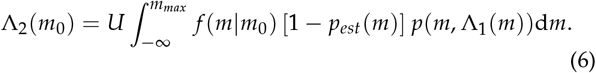

Following this logic, we can retrieve the probability of *k*-step rescue, for arbitrary *k* ≥ 2, using the recursion

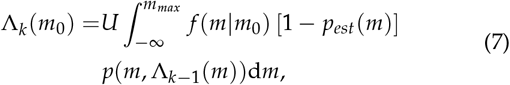

with the initial condition given by Equation 5.

### Approximating the probability of 2-step rescue

The probability of 2-step rescue is given by Equation 2 with *p*_0_ = *p*(*m*_0_, Λ_2_(*m*_0_)) (Equations 3–6). We next develop some intuition by approximating this for different classes of single mutants.

First, note that when the growth rate of a first-step mutation is close enough to zero such that *m*^2^ << Λ_1_(*m*), we can approximate the probability that such a genotype leads to rescue before itself going extinct, *p*(*m*, Λ_1_(*m*)), using a Taylor series, as 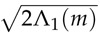 (c.f. equation A.4b in Iwasa *et al.* 2004a, see also File S1). We can also derive this result heuristically by considering the probability that a lineage will persist long enough that it will incur a secondary rescue mutation. As shown in the Appendix (see Mutant lineage dynamics), while *t* < 1/|*m*| a mutant lineage with growth rate *m* that is destined for extinction persists for *t* generations with probability ~ 2/*t* (Equation 21) and in generation *t* since it has arisen has ~ *t*/2 individuals (Equation 22). Thus, while *T* < 1/|*m*| a mutant lineage that persists for *T* generations will have produced a cumulative number ~ *T*^2^/4 individuals. Such lineages will then lead to 2-step rescue with probability ~ Λ_1_(*m*)*T*^2^/4 until this approaches 1, near 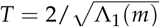. Since the probability of rescue increases like *T*^2^ while the probability of persisting to time *T* declines only like 1/*T*, most rescue events will be the result of rare long-lived single mutant genotypes. Considering only the most long-lived genotypes, the probability that a first-step mutation leads to rescue is then the probability that it survives long enough to almost surely rescue, i.e., for 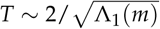 generations. Since the probability of such a long-lived lineage is 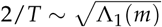, this heuristic result agrees with our Taylor series approximation of Equation 5. Thus, for first-step mutants with growth rates satisfying 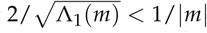, implying *m*^2^ << Λ_1_(*m*), which occur with probability 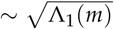, persistence is long enough to almost certainly ensure rescue. This same reasoning has been used to explain why the probability that a neutral mutation segregates long enough to produce a second mutation is 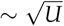 in a population of constant size (Weissman *et al.* 2009).

At the other extreme, when the growth rate of a first-step mutation is far enough from zero such that *m*^2^ >> Λ_1_(*m*), we can approximate *p*(*m*, Λ_1_(*m*)), again using a Taylor series, with Λ_1_(*m*)/|*m*| (c.f. equation A.4c in Iwasa *et al.* 2004a, see also File S1). Conditioned on extinction such genotypes cannot persist long enough to almost surely lead to 2-step rescue. Instead, we expect such mutations to persist for at most ~ 1/|*m*| generations (Equation 21) with a lineage size of ~ 1 individual per generation (Equation 22), and thus produce a cumulative total of ~ 1/|*m*| individuals. The probability of 2-step rescue from such a first-step mutation is therefore Λ_1_(*m*)/|*m*|, and again this heuristic argument matches our Taylor series approach. This same reasoning explains why a rare mutant genotype with selection coefficient |*s*| >> 0 in a constant population size model is expected to have a cumulative number of ~ 1/|*s*| descendants, given it eventually goes extinct (Weissman *et al.* 2009).

The transitions between these two regimes occur when 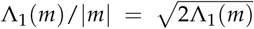, i.e., when 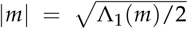. We call single mutants with growth rates 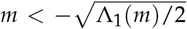 “sufficiently subcritical”, those with 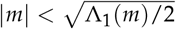 “sufficiently critical”, and those with 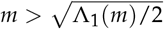 “sufficiently supercritical”. Given that *U* and thus Λ_1_(*m*) will generally be small, *m* will also be small at these transition points, meaning we can approximate the transition points as 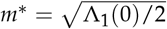 and −*m*^∗^. We then have an approximation for the rate of 2-step rescue,

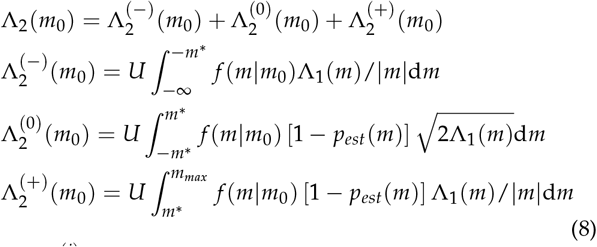

where 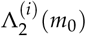 is the rate of 2-step rescue through sufficiently subcritical first-step mutants (*i* = “ − ”), sufficiently critical first-step mutants (*i* = 0), or sufficiently supercritical first-step mutants (*i* = “ + ”). A schematic depicting the 1- and 2-step genetic paths to rescue is given in Figure 4.

### Closed-form approximation for critical 2-step rescue

When *U* is small *m*^∗^ is also small, allowing us to use *m* = 0 within the integrand of 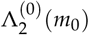, which spans a range, [−*m*^∗^, *m*^∗^], of width 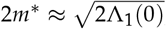, giving

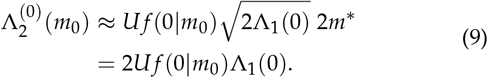

We can then approximate Λ_1_(*m*) with 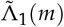 (Equation 19) and take *m* → 0 (Equation 20), giving a closed form approximation for the rate of 2-step rescue through critical single mutants in Fisher’s geometric model,

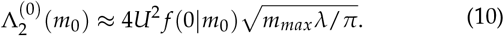

This well approximates numerical integration of 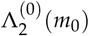 (Equation 8; see Figure 5 and File S1). In general, it will perform better when the critical zone, and thus 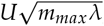, becomes smaller.

To get a better understanding of how the rate of 2-step critical rescue depends on the underlying parameters of Fisher’s geometric model, we approximate *f*(*m*|*m*_0_), assuming that the distance from the wildtype to the optimal phenotype is large relative to the distribution of mutations (i.e., *ρ*_*max*_ = *m*_*max*_/*λ* is large), and convert this to a distribution over 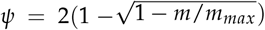, a convenient rescaling (for details see File S1 and Anciaux *et al.* 2018). Evaluating this at *m* = 0 gives

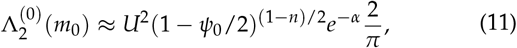

where 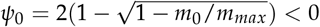 and 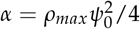.

### Closed-form approximations for non-critical 2-step rescue

We can also approximate Λ_1_(*m*) in 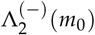 and 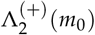 with 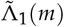 (Equation 19), leaving us with just one integral over the growth rates of the first-step mutations. We then replace *f* (*m*|*m*_0_) with its approximate distribution over *ψ* as above.

In the case of subcritical rescue we can then make two contrasting approximations (see File S1 for details). First, when the *ψ* (and thus *m*) that contribute most are close enough to zero (meaning maladaptation is not too large relative to mutational variance) and we ignore mutations that are less fit than the wildtype, we find the rate of subcritical 2-step rescue is roughly

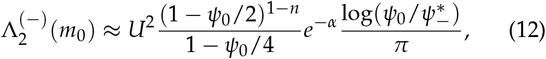

where 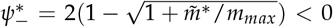 and 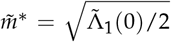 (Equation 20). Second, when the mutational variance, *λ*, is very small relative to maladaptation, implying that mutants far from *m* = 0 substantially contribute, we find the rate of subcritical 2-step rescue to be nearly

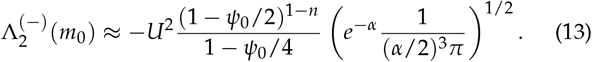

These two approximations do well compared with numerical integration of 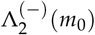 (Equation 8; see Figure 5 and File S1). As expected, we find that Equation 13 does better under fast wild-type decline while Equation 12 does better when the wildtype is declining more slowly.

For supercritical 2-step rescue, only first-step mutants with growth rates near *m*^∗^ will contribute (larger *m* will rescue themselves and are also less likely to arise by mutation), and so we can capture the entire distribution with a small *m* approximation (following the same approach that led to Equation 12). As shown in File S1, this approximation works well for sufficiently small first-step mutant growth rates, 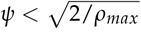, beyond which the rate of 2-step rescue through such first-step mutants falls off very quickly due to a lack of mutational input. Thus, considering only supercritical single mutants with scaled growth rate less than 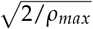, our approximation is

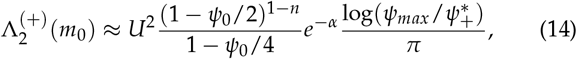

with 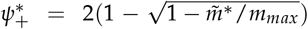 and 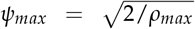. This approximation tends to provide a slight overestimate of 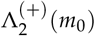 (Equation 8; see Figure 5 and File S1).

### Comparing 2-step regimes

These rough but simple closed-form approximations (Equations 11–14) show that, while the contribution of critical mutants to 2-step rescue scales with *U*^2^, the contribution of non-critical single mutants scales at a rate less than *U*^2^ (Figure 5B) due to a decrease in 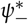 (decreasing the range of subcritical mutants) and an increase in 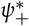 (decreasing the range of supercritical mutants) with *U*. This difference in scaling with *U* is stronger when the wildtype is not very maladapted relative to the mutational variance, i.e., when Equation 12 is the better approximation for subcritical rescue. The approximations also show that when initial maladaptation is small, the ratio of supercritical to subcritical contributions (Equation 12 divided by 14) primarily depends on the range of growth rates included in each regime, while with larger initial maladaptation this ratio (Equation 13 divided by 14) begins to depend more strongly on initial maladaptation and mutational variance (*α*). The effect of maladaptation and mutation rate on the relative contributions of each regime is shown in Figure 5.

### The distribution of growth rates among rescue genotypes

We next explore the distribution of growth rates among rescue genotypes, i.e., the distribution of growth rates that we expect to observe among the survivors across many replicates.

We begin with 1-step rescue. The rate of 1-step rescue by genotypes with growth rate *m* is simply *Uf* (*m*|*m*_0_)*p*_*est*_(*m*). Dividing this by the rate of 1-step rescue through any *m* (Equation 5) gives the distribution of growth rates among the survivors

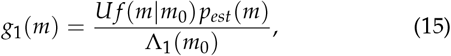

where the mutation rate, *U*, cancels out. This distribution is shown in blue in Figure 6. The distribution has a mode at small but positive *m* as a result of two conflicting processes: smaller growth rates are more likely to arise from a declining wildtype but larger growth rates are more likely to establish given they arise. As the rate of wildtype decline increases, the former process exerts more influence, causing the mode to move towards zero and reducing the variance.

We can also give a simple, nearly closed-form approximation here using the same approach taken to reach Equation 19. On the *ψ* scale, the distribution of effects among 1-step rescue mutations is

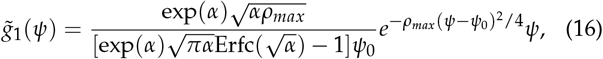

implying the *ψ* are distributed like a normal truncated below *ψ* = 0 and weighted by *ψ*. This often provides a very good approximation (see dashed blue curves in Figure 6).

In 2-step rescue, the rate of rescue by double mutants with growth rate *m*_2_ is given by Equation 6 with Λ_1_(*m*) replaced by *Uf*(*m*_2_|*m*)*p*_*est*_(*m*_2_). Normalizing gives the distribution of growth rates among the double mutant genotypes that rescue the population

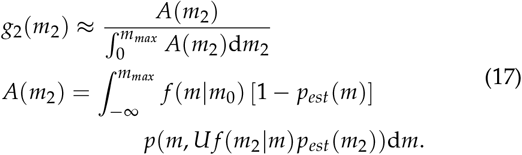

This distribution, *g*_2_(*m*), is shown in red in Figure 6. Because the first-step mutants contributing to 2-step rescue tend to be nearer the optimum than the wildtype, this allows them to produce double mutant rescue genotypes with higher growth rates than in 1-step rescue (as seen by comparing the mode between blue and red curves in Figure 6). The fact that these first-step mutants are closer to the optimum also allows for a greater variance in the growth rates of rescue genotypes than in 1-step rescue. Thus the 2-step distribution maintains a more similar mode and variance across wildtype decline rates than the 1-step distribution. Note that because *g*_2_(*m*_2_) depends on *U* the buffering effect of first-step mutants depends on the mutation rate (see The distribution of growth rates among rescue intermediates below for more discussion).

### The distribution of growth rates among rescue intermediates

Finally, our analyses above readily allow us to explore the distribution of first-step mutant growth rates that contribute to 2-step rescue. Analogously to Equation 15, we drop the integral in Λ_2_(*m*_0_) (Equation 6) and normalize, giving

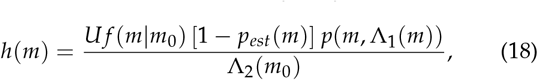

where the first *U* cancels but the *U* within Λ_1_(*m*) does not. This distribution is shown in black in Figure 7. At slow wildtype decline rates the overwhelming majority of 2-step rescue events arise from first-step mutants with growth rates near 0. As indicated by Equation 8, the contribution of first-step mutants with growth rate *m* declines as ~ 1/|*m*| as *m* departs from zero, due to shorter persistence times given eventual extinction. As wildtype growth rate declines, the relative importance of mutational input, *f*(*m*|*m*_0_), grows, causing the distribution to flatten and first-step mutants with substantially negative growth rates begin to contribute (compare panels in Figure 7; see also Figure 5A). Decreasing the mutation rate disproportionately increases the contribution of first-step mutants with growth rates near zero (while simultaneously shrinking the range of growth rates that are sufficiently critical; Figure 5B) making the distribution of first-step mutant growth rates contributing to 2-step rescue more sharply peaked around *m* = 0 (Figure S2). Correspondingly, with a higher mutation rate a greater proportion of the contributing single mutants have substantially negative growth rates.

## Discussion

Here we have explored the probability and genetic basis of evolutionary rescue by multiple mutations on a simple fitness landscape. We find that rescue by multiple mutations can be the most likely path to persistence under high mutation rates or when the population is initially very maladapted. Under these scenarios, intermediate genotypes that are declining less quickly provide a ‘springboard’ from which rescue genotypes emerge. In 2-step rescue these springboard single mutants come from one of three regimes: those that have growth rates near enough to zero (“sufficiently critical”) that rescue is most likely when a mutation persists for an unusually long period of time and grows to an unusually large subpopulation size, and those with growth rates that are either negative or positive enough (“sufficiently subcritical” or “sufficiently supercritical”, respectively) to restrict persistence times and subpopulation sizes, conditioned upon the loss of the first mutation in the absence of a second, rescuing mutation. The relative contribution of each regime shifts with initial maladaptation and mutation rate; rare mutations that can occasionally reach unusually large subpopulation sizes play a larger role when the population is not severely maladapted (e.g., Figure 7A) or mutation rate is high (e.g., Figure S2C). In contrast, when populations are initially very maladapted (e.g., Figure 7C), most first-step mutations are themselves also very maladapted and thus restricted in the subpopulation sizes they are expected to reach before being lost. All three regimes help to maintain the variance in the distribution of fitness effects among rescue genotypes as initial maladaptation increases; meanwhile, in 1-step rescue the variance declines due to ever more extreme sampling of the tail of the mutational distribution (compare blue and red curves in Figure 6).

Our prediction, that rescue by more *de novo* mutations can be more likely than rescue by fewer, is novel. In previous models (e.g., Antia *et al.* 2003; Iwasa *et al.* 2004a; Alexander and Day 2010) the general conclusion has been that, since the probability of rescue scales with *U*^*k*^ (where *U* is the mutation rate and *k* is the minimum number of mutations required for rescue), the probability of rescue declines with the number of mutations. This assumes, however, that the probability of a mutation occurring, *U*, is the limiting factor. Here we have shown that when the probability of a beneficial mutation arising declines with its selective advantage, the probability of sampling once from the extreme tail of the DFE can be lower than sampling multiple mutations closer to the bulk of the DFE, so that rescue via multiple mutations can become the dominant path. Rescue by multiple mutations may also be more likely with standing genetic variation, as small-effect intermediate mutations may segregate at higher frequencies than large-effect rescue mutations before the environmental change (and also decline less quickly than the wildtype following environmental change); this is especially true with recombination, where rescue genotypes can arise from segregating intermediate mutations without mutation (Uecker and Hermisson 2016).

How often rescue arises as a result of multiple mutations is an open question. It is clear that more than one mutation can contribute to adaptation to near-lethal stress, but experiments are often designed to avoid extinction (reviewed in Cowen *et al.* 2002) and therefore greatly expand the scope for multiple mutations to arise on a single genotype. A few exceptions provide some insight. For example, populations of *Saccharomyces cervisae* that survived high concentrations of copper acquired multiple mutations (Gerstein *et al.* 2015) – in fact the authors argue for the ‘springboard effect’ discussed above, where first-step mutations prolong persistence and thereby allow further mutations to arise. In *Pseudomonas flourescens*, fluctuation tests with nalidixic acid showed that nearly a third of the most resistant surviving strains were double mutants (Bataillon *et al.* 2011), which were able to tolerate 10x higher drug concentrations than single mutants, suggesting 2-step rescue might dominate at high drug concentrations. While suggestive, it is unclear if our prediction – that rescue takes more mutational steps with greater initial maladaptation – holds true generally. Verification will require more experiments that allow extinction and uncover the genetic basis of adaptation at different severities of environmental change (e.g., drug concentration).

In describing the genetic basis of adaptation in populations of constant size, Orr (1998) showed that the mean phenotypic displacement towards the optimum scales roughly linearly with initial displacement. Converting phenotype to fitness, this implies that the mean fitness effect of fixed mutations (*s* = *m* − *m*_0_) increases exponentially as initial Malthusian fitness (*m*_0_) declines (i.e., *s* ~ exp(−*m*_0_)), which is a roughly linear increase when initial fitness is small (|*m*_0_| << 1). Here we see that, under 1-step rescue, the mean fitness effect also increases roughly linearly as the initial growth rate declines (see horizontal blue lines in Figure 6). However, the rate of this linear increase in fitness effect is much larger under rescue than in a population of constant size (compare blue and grey horizontal lines in Figure 6), where declines in wildtype fitness not only allow larger mutations to be beneficial but also require larger mutations for persistence. Thus the race between extinction and adaptation during evolutionary rescue is expected to produce a genetic basis of adaptation with fewer mutations of larger effect.

While under 1-step rescue the fitness effect of the first mutation increases roughly linearly as wildtype fitness declines, most rescue events will be 2-step for wildtype fitnesses below some value (e.g., at *m*_0_ ≈ −0.25 in Figure 3; this threshold value of *m*_0_ increases with mutation rate, Figure S1). At this junction the effect size of the first mutation will no longer increase as quickly (and potentially even decrease), as it switches from a rescue mutant to an intermediate mutant whose expected fitness begins to decline substantially with the fitness of the wildtype (Figure 7). Thus as rescue switches from dominantly *k*-step to dominantly (*k* + 1)-step the genetic basis of adaptation becomes more diffuse, with each mutation having a smaller individual fitness effect as the contributing fitness effects spread over more loci. In the limit of large *k* (due to large initial maladaptation or high mutation rates), the genetic basis of adaptation should at some point converge to many loci with small effect, as would also be expected in a population of constant size. Indeed, at very high mutation rates the rate of adaptation (the change in mean fitness) is the same under rescue as it is in populations of constant size (Anciaux *et al.* 2019), implying that the genetic basis of adaptation no longer depends on demography. It is therefore at intermediate levels of initial maladaptation and low mutation rates, where rescue primarily occurs from a few large effect mutations, that the race between adaptation and persistence is predicted to have the largest effect on the genetic basis of adaptation.

Fluctuation tests (Luria and Delbrück 1943) provide a means to generate random mutations and then isolate potential rescue genotypes (typically assumed to be 1-step only), whose growth rates can be measured under the selective conditions. These experiments are designed such that there is substantial standing genetic variation at the time of exposure to the selective conditions, which should increase the contributions of mutations with small growth rates (Orr and Betancourt 2001), although these could be outcompeted by mutations with higher growth rates and/or be under-sampled. Regardless, consistent with our theory (Figure 6), the resulting growth rate distributions in both bacteria and yeast often find modes that are substantially greater than zero (as opposed to, say, an exponential distribution; Kassen and Bataillon 2006; MacLean and Buckling 2009; Gerstein *et al.* 2012; Lindsey *et al.* 2013; Gerstein *et al.* 2015). A number of these conform even more closely to our expected shape (Kassen and Bataillon 2006; Gerstein *et al.* 2015) while the others appear to be substantially more clumped around the mode, perhaps due to a very restricted number of possible rescue mutations in any one circumstance, the size of the experiment, or the way in which growth rates are measured. Finally, Gerstein *et al.* (2015) not only provide the distribution of growth rates among rescue genotypes, but also the growth rates of individual mutations that compose multi-step rescue genotypes. In four lines where multiple mutations were detected and a segregation analysis performed, one mutation in each line was inferred to have a minor effect and the other mutation was an amplification of the copper metallothionein CUP with a major fitness effect. These results are consistent with the minor effect mutations being sub-critical mutations that provided a springboard for the larger CUP mutations.

Pinpointing the mutations responsible for adaptation is hampered by genetic hitchhiking, as beneficial alleles elevate the frequency of linked neutral and mildly deleterious alleles (Barton 2000). The problem is particularly severe under strong selection and low recombination, and therefore reaches an extreme in the case of evolutionary rescue in asexuals, especially if many neutral and deleterious mutations are segregating at the time of environmental change. To circumvent this, mutations that have risen to high frequency in multiple replicates are often introduced in a wildtype background, in isolation and sometimes also in combination with a small number of other common highfrequency mutations, and grown under the selective conditions (e.g., Jochumsen *et al.* 2016; Ono *et al.* 2017). As we have demonstrated above (e.g., Figure 7C), however, under multi-step rescue there may be no one mutation that individually confers growth in the selective conditions. Thus, a mutation that was essential for rescue may go undetected or be mistaken as a hitchhiker if the appropriate multiple-mutation genotypes are not tested. Unfortunately reverse engineering all combinations of mutations quickly becomes unwieldy as the number of mutations grows, and thus this approach will not be practical under severe initial maladaptation and high mutation rates, where we predict rescue to occur by many mutations. Interestingly, our simulations show that the population dynamics themselves may help differentiate how many mutations contribute to rescue (e.g., V- vs. U-shaped log-trajectories; Figures 1 and 2), and fitting models of *k*-step rescue could produce estimates for the growth rates of the *k* genotypes.

Environmental change often selects for mutator alleles, which elevate the rate at which beneficial alleles arise and subsequently increase in frequency with them (Tenaillon *et al.* 2001). When beneficial alleles are required for persistence, as in evolutionary rescue, mutator alleles can reach very high frequencies or rapidly fix (e.g., Mao *et al.* 1997). Consistent with this, mutator alleles are often associated with antibiotic resistance in clinical isolates (see examples in Bell 2017). Further, the more beneficial mutations available the larger the advantage of a mutator allele; for a mutator that increases the mutation rate *m*-fold, its relative contribution to the production of *n* beneficial mutations scales as *m*^*n*^ (Tenaillon *et al.* 1999). Thus, conditions that cause multi-step rescue to be more likely than 1-step rescue should also impose stronger selection for mutator alleles. There are a number of examples where lineages with higher mutation rates acquired multiple mutations and persisted at higher doses of antibiotics (Couce *et al.* 2015; San Millan *et al.* 2017). The number of mutations required for persistence is, however, often unknown, making it difficult to compare situations where rescue requires different numbers of mutations. Experiments with a combination of drugs may provide a glimpse; for instance, *Escherichia coli* populations only evolved resistance to a combination of two drugs (presumably through the well-known mutations specific to each drug) when mutators were present, despite the fact that mutators were not required for resistance to either drug in isolation (Gifford *et al.* 2019). In cases where we have less information on the genetic basis of resistance, our model suggests that mutators will be more advantageous when initial maladaptation is severe (e.g., higher drug concentrations or a larger number of drugs), as rescue will then be dominated by genetic paths with more mutational steps.

Here we have investigated the genetic basis of evolutionary rescue in an asexual population that is initially genetically uniform. Extending this work to allow for recombination and standing genetic variation at the time of environmental change – as expected for many natural populations – would be valuable. The effect of standing genetic variation on the probability of 1-step rescue is relatively straight-forward to incorporate, depending only on the expected number of rescue mutations initially present and their mean establishment probability (Martin *et al.* 2013). In the case of the fluctuation tests discussed above, where mutations accumulated in the short interval before the onset of selection are assumed to be relatively neutral, the effect of standing genetic variance on 1-step rescue might be incorporated by a simple rescaling of *N*_0_, to account for the additional mutants present in the standing variation. When considering longer periods of time in populations that are not rapidly expanding, mutation-selection balance may be reached before the onset of selection. In this case the probability of 1-step rescue from standing genetic variance in Fisher’s geometric model was given by Anciaux *et al.* (2018), whose equations 3 and 5 immediately give the distribution of fitness effects among those that rescue. Allowing these standing genetic variants to be springboards to multi-step rescue will help clarify the role of standing genetic variation on the genetic basis of rescue more generally. Recombination can help combine such springboard mutations into rescue genotypes but will also break these combinations apart, as demonstrated in a 2-locus 2-allele model of rescue (Uecker and Hermisson 2016). How recombination affects the genetic basis of evolutionary rescue when more loci can potentially contribute remains to be seen. Also left unexplored is the effect of density-dependent fitness; for example, competition may reduce mutant growth rates and thereby increase the size of mutations that are required for rescue, especially when the wildtype declines slowly. Combining density-dependence and standing genetic variance is known to create complex dynamics in a 1-locus 2-allele model of rescue (Uecker *et al.* 2014), and adding more potential genotypes is sure to add yet more complexity.

Many of our simple closed-form results rely upon knowing the distribution of mutant growth rates (Equation 1), which arises from the assumption that mutant phenotypes are normally distributed about their ancestor and Malthusian fitness is a quadratic, on some scaled phenotypic axes. It is clear that deviations from these assumptions will, at least quantitatively, affect our results. For instance, mutant phenotype distributions with truncated or fat tails are likely to lead to smaller or larger mutational steps, respectively, with downstream effects on the probability of rescue, the number of contributing mutations, and the resulting DFEs. As a preliminary investigation of this prediction, we have performed simulations with mutant phenotype distributions having the same expectation and covariances as assumed above under normality, but with truncated (platykurtic) or fat (leptokurtic) tails (Figure S3A). While our qualitative results above hold, the probability of rescue declines slower with wildtype maladaptation when the mutational distribution has fatter tails (compare dotted and solid black in Figure S3C). Fatter tails also reduce the number of mutations contributing to rescue (e.g., 1-step rescue dominates for all wildtype decline rates in Figure S3C). Finally, fatter tails cause the distributions of rescue genotype growth rates following 1- and 2-step rescue to have more variance and become more similar to one another (Figure S4B) and also tend to increase the contribution of supercritical single mutants in 2-step rescue (Figure S5). All told, the genetic basis of rescue is expected to consist of fewer mutations of larger effect, with less consistent effect sizes across replicate populations, as the tails of the mutant phenotype distribution become fatter.

In the numerical examples above we have not varied the number of scaled phenotypic axes, *n*, i.e., the dimensionality of the phenotypic landscape (although the analytical results apply for arbitrary *n*). Because increasing the number of dimensions changes the distribution of fitness effects, and in particular decreases the proportion of mutations that are beneficial (Fisher 1930), this may have cascading influences on our results. As shown in Anciaux *et al.* (2018), the probability of 1-step rescue by *de novo* mutation declines with dimensionality, and is only weakly dependent on dimensionality when initial maladaptation is small (such that Λ_1_(*m*_0_) ≈ −*m*_0_*Ug*(*α*), Equation 19). Here we show that the distribution of fitness effects among 1-step rescue mutants is nearly independent of dimensionality for any degree of initial maladaptation (Equation 16 and the blue curves in Figure S6B). Further, as seen by comparing Equations 11–14 to Equation 19, the probability of 2-step rescue depends on dimensionality much like 1-step rescue does, suggesting that while increasing dimensionality may decrease the probability of rescue it may have little effect on the number of steps rescue tends to take. This is demonstrated more generally in Figure S6A, where an order of magnitude increase in the number of dimensions decreases the probability of rescue by roughly an order of magnitude but has little effect on the relative rates of 1-, 2-, 3-, and 4-step rescue. Finally, Figure S6B-C shows that dimensionality has very little effect on the distribution of fitness effects among 2-step rescue genotypes (Equation 17) and among first step mutants leading to 2-step rescue (Equation 18). To conclude, while the probability of rescue declines with the complexity of the organism and its environment, the genetic basis of rescue is expected to be relatively invariant across complexity, as with the genetic basis of adaptation in populations of constant size (Orr 1998, see also gray curves in Figure S6B,C).

In the numerical examples above we have also focused on a particular value of mutational variance, *λ*. Clearly, since rescue relies on mutations of large effect, decreasing *λ* should decrease the probability of rescue, much like decreasing the mutation rate, *U*, does (Figure S1). While our analysis (Equations 19 and 11-14) and numerical results (see File S1) show that this is true, we find that *λ* and *U* have very different effects on the genetic basis of rescue (File S1). In particular, given a similar effect on the total probability of rescue, decreasing *U* generally restricts rescue to fewer mutational steps while decreasing *λ* forces rescue to occur by more mutations. Further, the distribution of fitness effects of mutations contributing to rescue is nearly independent of *U* but a decrease in *λ* strongly reduces the mode of the DFE. This demonstrates that populations with similar probabilities of rescue can vary greatly in the way they achieve it genetically.

## Acknowledgements

We would like to thank the Otto and Doebeli labs for helpful feedback at various stages, Ophélie Ronce and Thomas Lenormand for their hospitality and valuable input at the beginning of this project, and Mike Whitlock, Amy Angert, Luis-Miguel Chevin, and Joachim Hermisson for constructive criticism on previous versions of the manuscript. Funding provided by the National Science and Engineering Research Council (CGS-D 6564 to M.M.O., RGPIN-2016-03711 to S.P.O.), the University of British Columbia, Banting, and the University of California - Davis (fellowships to M.M.O.), the National Institute of General Medical Sciences of the National Institutes of Health (NIH R01 GM108779 to Graham Coop), the Agence Nationale de la Recherche (ANR-18-CE45-0019 “RESISTE” to G.M.), and the Centre Méditerranéen Environment et Biodiversité (“BACTPHI” to G.M.).

## Appendix

## Approximating the probability of 1-step rescue

The probability of 1-step rescue in this model has been derived by Anciaux *et al.* (2018). As replicated in File S1 and given by their equation 7, when *ρ*_*max*_ = *m*_*max*_/*λ* is large a simple, nearly closed-form approximation is

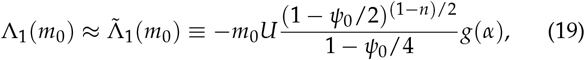

where 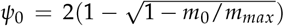, 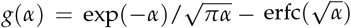, and 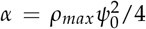, with erfc(.) the complimentary error function. When the wildtype declines slowly *m*_0_ and thus *ψ*_0_ is small and Λ_1_(*m*_0_) ≈ *Ug*(*α*). In the limit *m*_0_ → 0, Equation 19 becomes

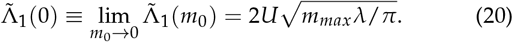

## Mutant lineage dynamics

Here we follow the lead of Weissman *et al.* (2010) and Uecker and Hermisson (2016) in approximating our discrete-time process with a continuous-time branching process (see chapter 6 in Allen 2010). Consider a birth-death process, where individuals give birth at rate *b* and die at rate *d*. One can then obtain the probability generating function for the number of individuals at a given time, *n*(*t*), given the initial number, *n*(0). We are primarily interested in new mutant lineages, *n*(0) = 1. The generating function then allows us to calculate the probability that a lineage persists at least until time *t* and the distribution of *n*(*t*) given it does so (see below).

To convert between birth and death rates and our compound Malthusian parameter we follow Uecker and Hermisson (2016) in equally distributing the growth rate *m* between birth and death, *b* = (1 + *m*)/2 and *d* = (1 − *m*)/2, such that *m* = *b* − *d* and the continuous-time process exhibits the same amount of drift as the discrete time process (and matches discrete-time simulations well; Uecker *et al.* 2014). We can now report the necessary results in terms of *m* (assuming |*m*| < 1).

Denoting the extinction time as *T*, the probability a mutant with growth rate *m* persists until time *t* is approximately (see File S1 for derivation)

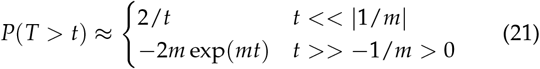

As pointed out in Weissman *et al.* (2010) (whose equation A2 differs from Equation 21 by a factor of 2 because they have *b* + *d* = 2), the distribution of persistence times has a long tail (like 1/*t*) until being cut off (declining exponentially) at *t* = −1/*m*.

Given a lineage persists until *t*, the distribution of *n*(*t*) is roughly (see File S1 for derivation)

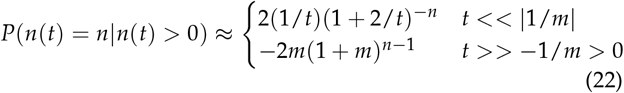

As pointed out in Weissman *et al.* (2010) (whose equation A3 only differs from Equation 22 by constants), the distribution of *n*(*t*) is approximately geometric for small or large *t*, implying *n*(*t*) is very unlikely to be greater than the minimum of *t* and −1/*m*.

---

**Figure S1.**
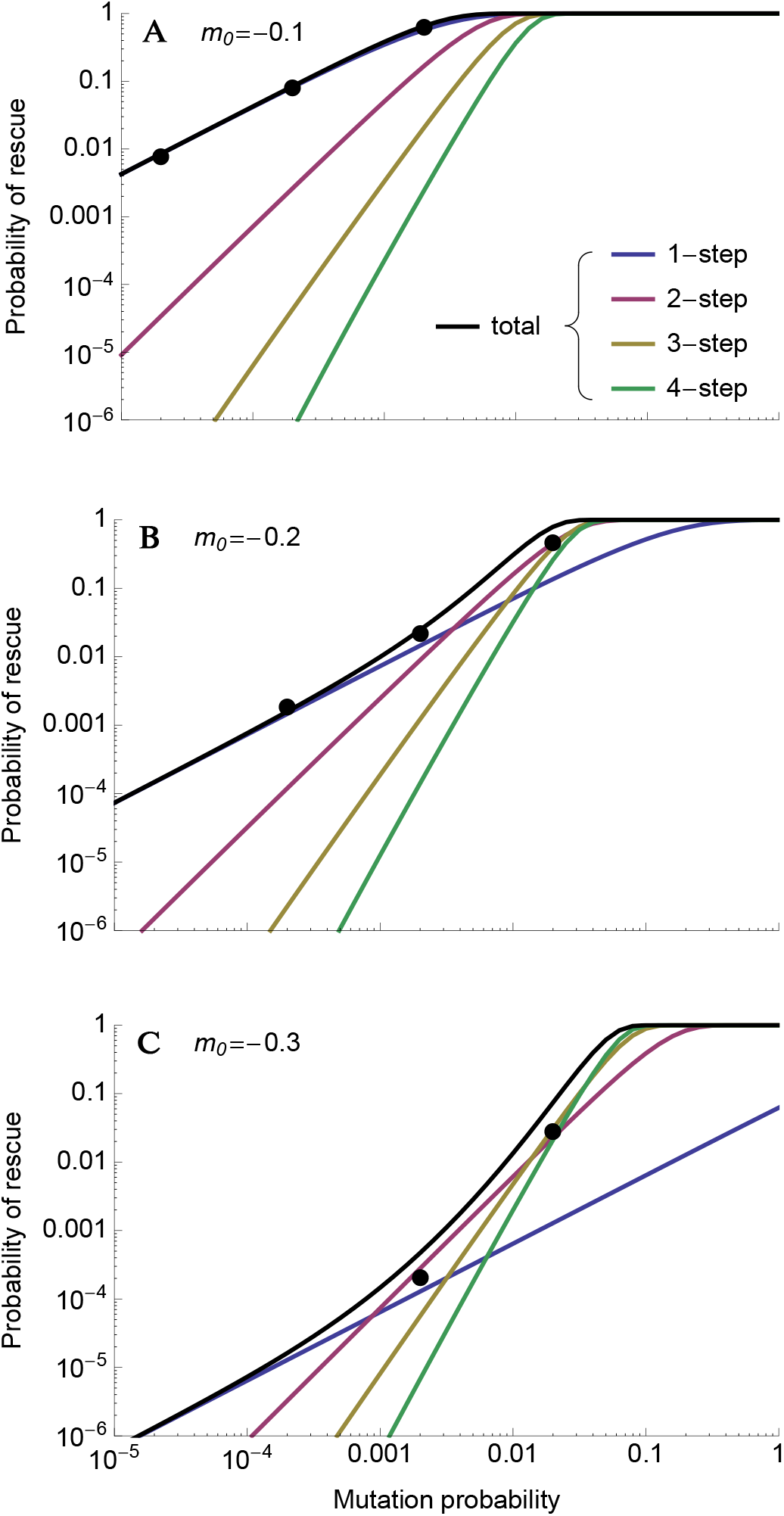
The probability of rescue as a function of mutation rate for three different levels of initial maladaptation. See Figure 3 for details. Other parameters: *n* = 4, *λ* = 0.005, *m*_*max*_ = 0.5, *N*_0_ = 10^4^.

**Figure S2.**
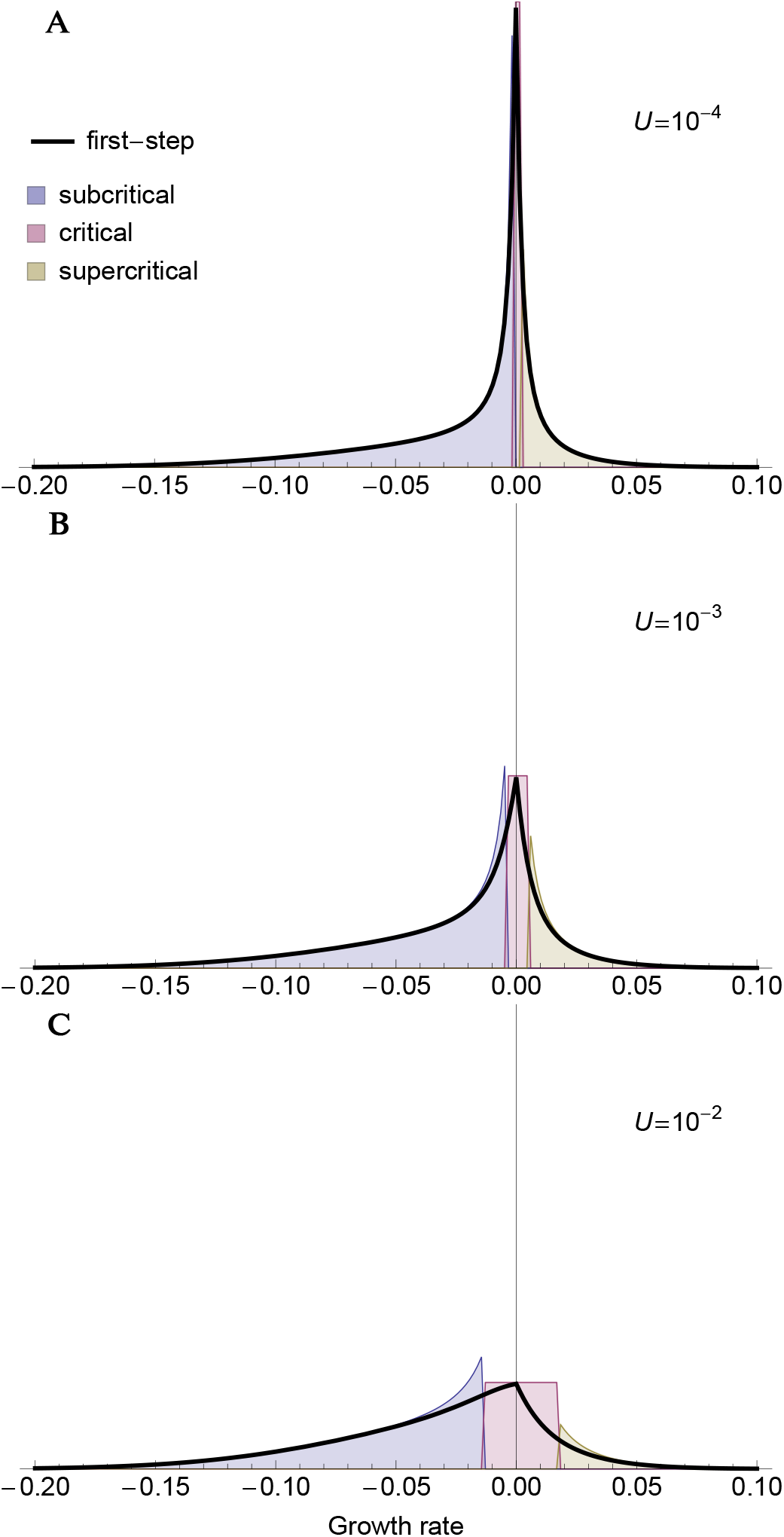
The distribution of first-step mutant growth rates given 2-step rescue under three mutation rates. See Figure 7 for details. Parameters: *n* = 4, *λ* = 0.005, *m*_*max*_ = 0.5, *m*_0_ = −0.2.

**Figure S3.**
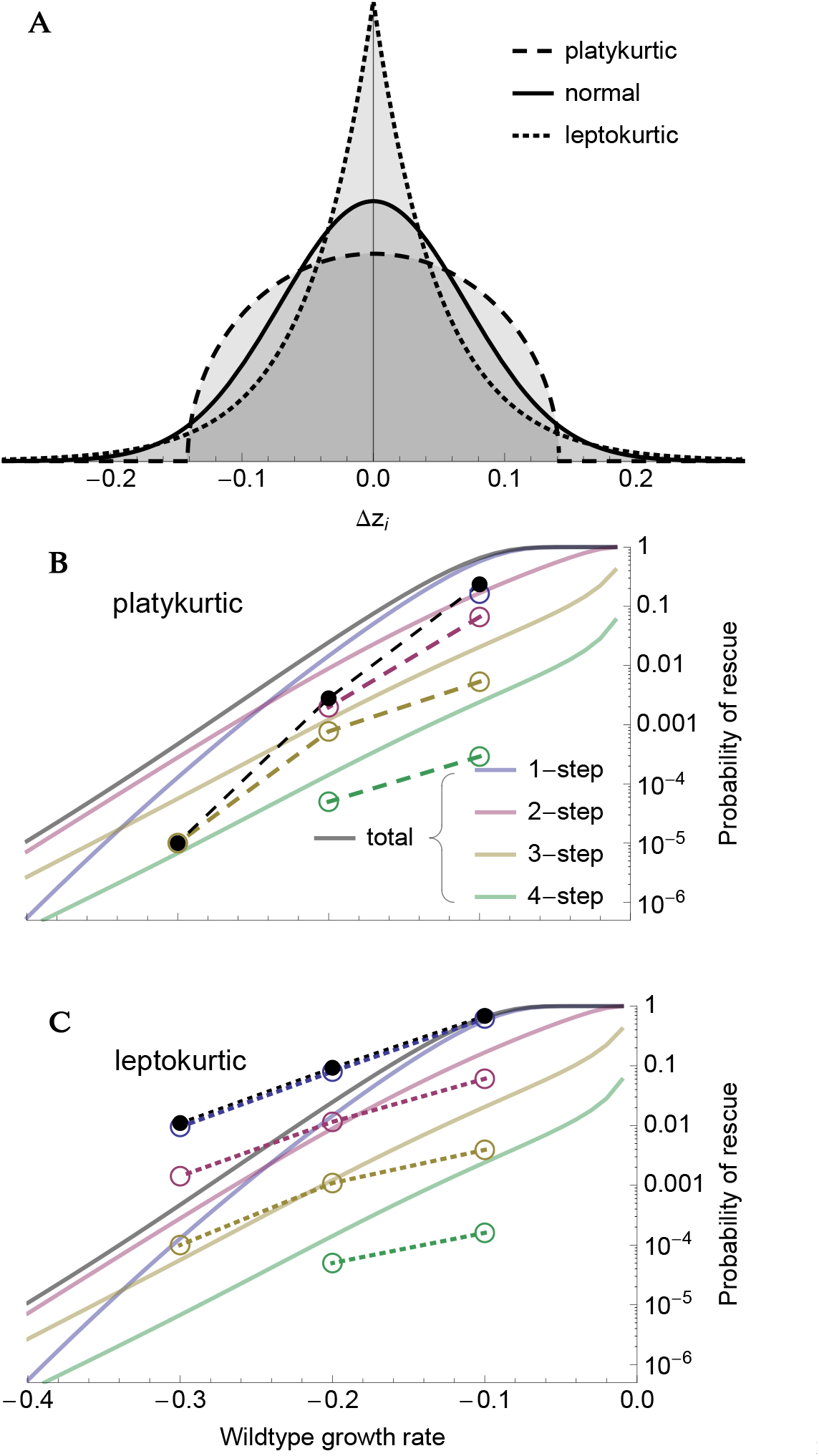
(**A**) One-dimensional slices of multidimensional platykurtic (dashed; semicircle), normal (solid; as used in main text), and leptokurtic (dotted; Laplace) mutational distributions with the same (co)variance but varying kurtosis. (**B**,**C**) The probability of 1-, 2-, 3-, or 4-step rescue with platykurtic and leptokurtic mutational distributions, respectively. The dots and broken lines represent simulation results (10^5^ replicates for each wildtype growth rate). The solid lines are the numerical results for the normal mutational distribution (as in Figure 3). Parameters: *N*_0_ = 10^4^, *U* = 2 × 10^−3^, *n* = 4, *λ* = 0.005, *m*_*max*_ = 0.5.

**Figure S4.**
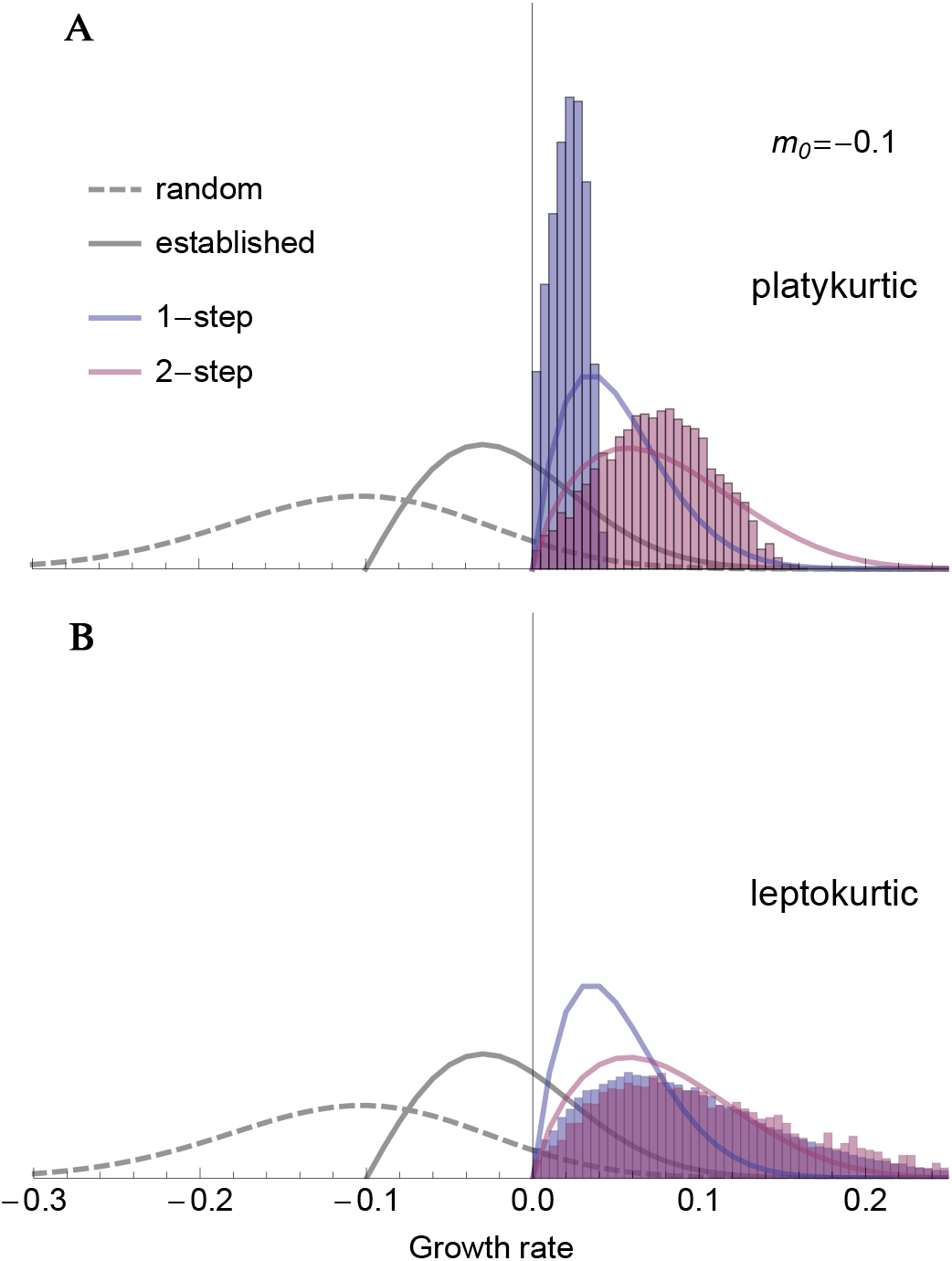
The distribution of growth rates among rescue genotypes under 1-step (blue) and 2-step (red) rescue with (**A**) platykurtic and (**B**) leptokurtic mutational distributions (see Figure S3A). The solid lines are predictions for a normal mutational distribution (as in Figure 6). The histograms show the distribution of growth rates among rescue genotypes observed across 10^5^ replicate simulations. Parameters: *N*_0_ = 10^4^, *U* = 2 × 10^−3^, *n* = 4, *λ* = 0.005, *m*_*max*_ = 0.5, *m*_0_ = −0.1.

**Figure S5.**
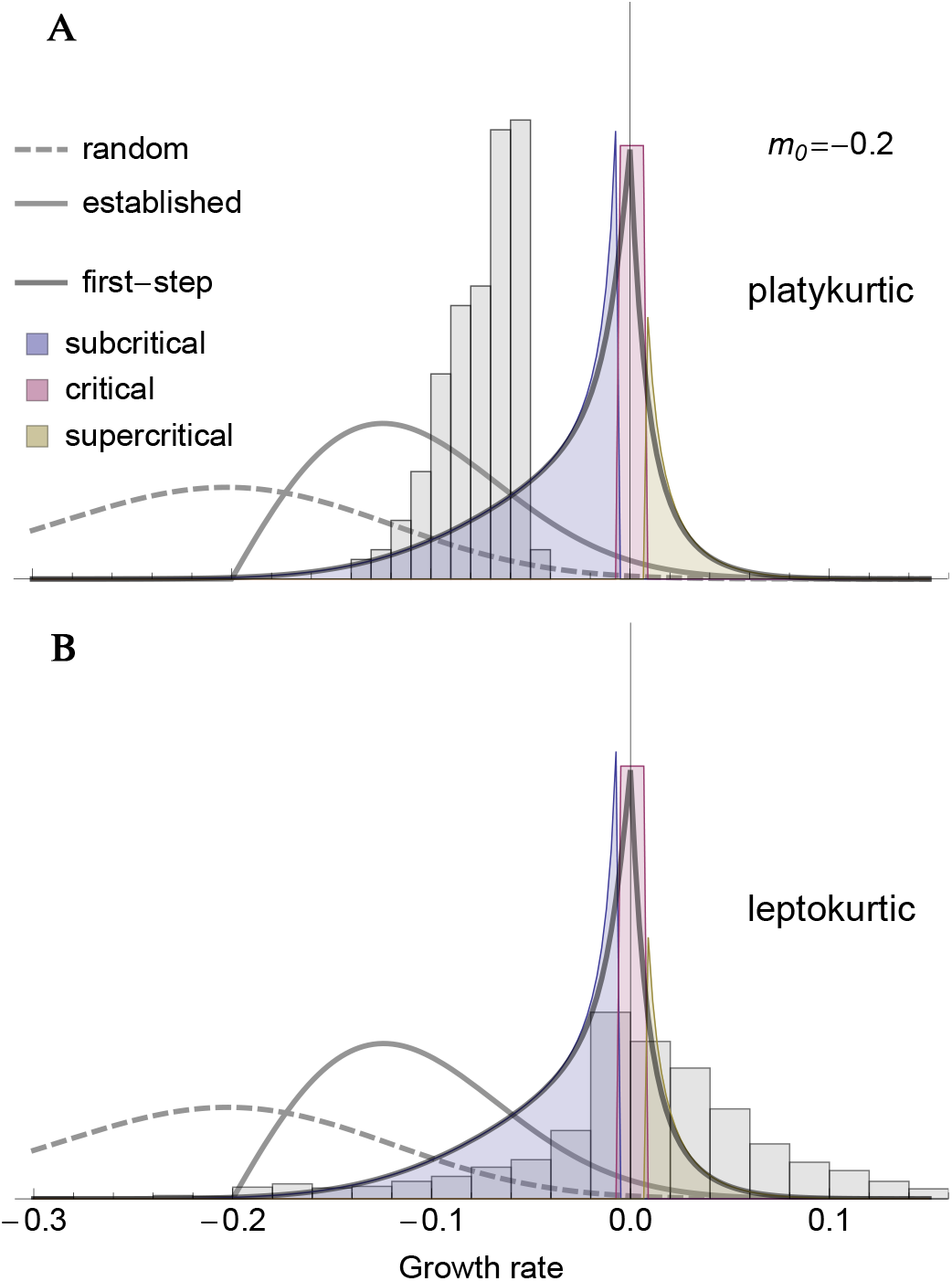
The distribution of growth rates among first-step mutations that lead to 2-step rescue with (**A**) platykurtic and (**B**) leptokurtic mutational distributions (see Figure S3A). The curves and shadings are predictions for a normal mutational distribution (as in Figure 7). The histograms show the distribution of growth rates observed across 10^5^ replicate simulations. Parameters: *N*_0_ = 10^4^, *U* = 2 × 10^−3^, *n* = 4, *λ* = 0.005, *m*_*max*_ = 0.5, *m*_0_ = −0.2.

**Figure S6.**
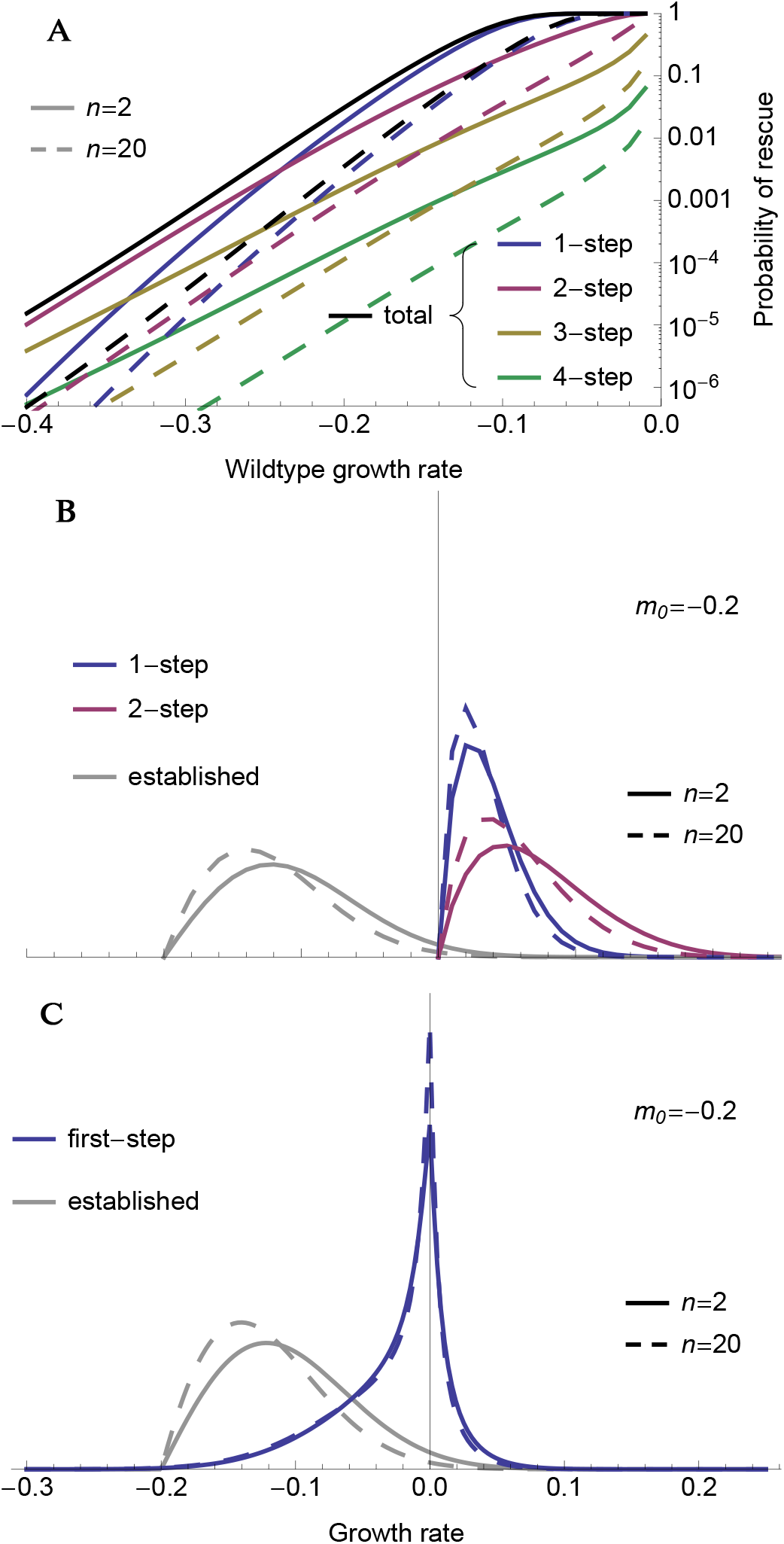
The effect of the number of phenotypic dimensions, *n*, on (**A**) the probability of *k*-step rescue, (**B**) the distribution of growth rates among rescue genotypes, and (**C**) the distribution of growth rates among first-step mutants that lead to 2-step rescue. Curves are numerical results, as in Figures 3, 6, and 7. Parameters: *N*_0_ = 10^4^, *U* = 2 × 10^−3^, *λ* = 0.005, *m*_*max*_ = 0.5.

